# Potent Protection Against H5N1 and H7N9 Influenza via Childhood Hemagglutinin Imprinting

**DOI:** 10.1101/061598

**Authors:** Katelyn M. Gostic, Monique Ambrose, Michae Worobey, James O. Lloyd-Smith

## Abstract

Two zoonotic influenza A viruses (IAV) of global concern, H5N1 and H7N9, exhibit puzzling differences in age distribution of human cases. Previous explanations cannot fully account for these patterns. We analyze data from all known human cases of H5N1 and H7N9 and show that an individual’s first IAV infection confers lifelong protection against severe disease from novel hemagglutinin (HA) subtypes of the same phylogenetic group. Statistical modeling reveals protective HA imprinting to be the crucial explanatory factor, providing 75% protection against severe infection and 80% protection against death for both H5N1 and H7N9. Our results enable us to predict age distributions of severe disease for future pandemics and to demonstrate that a novel strain’s pandemic potential increases yearly when a group-mismatched HA subtype dominates seasonal influenza circulation. These findings open new frontiers for rational pandemic risk assessment.

## Main Text

The spillover of novel influenza A virus (IAV) strains is a persistent threat to global public health. Current thinking assumes a central role for an immunologically naïve human population in the emergence of pandemic IAVs (*1*). H5N1 and H7N9 are particularly concerning avian-origin IAVs, each having caused hundreds of severe or fatal human cases (*2*). Despite commonalities in their reservoir hosts and epidemiology, these viruses show puzzling differences in age distribution of observed human cases (*2,3*). Explanations proposed to date, including possible protection against H5N1 from past exposure to the neuraminidase of H1N1 (*4,5*) and human behavioral factors (such as age bias in exposure to infected poultry) (*6–8*), cannot fully explain these opposing patterns of severe disease and mortality. Another idea is that severity of H5N1 and H7N9 differs by age class, leading to case ascertainment bias (*2*), but no explanatory mechanism has been proposed.

The key determinants for IAV susceptibility are the virus’s two surface glycoproteins, hemagglutinin (HA) and neuraminidase (NA), where different numbered subtypes canonically indicate no cross-immunity. However, recent experiments have revealed that broadly cross-protective immune responses can provide immunity between different HA subtypes, typically within the same phylogenetic group (*9–15*). H5 belongs to HA group 1 (which also includes H1 and H2), while H7 is in group 2 (which includes H3). We hypothesized that the 1968 pandemic, which marked the transition from an era of group 1 HA circulation (1918-1968) to an era dominated by a group 2 HA virus (1968-present) (Fig. 1A), caused a major shift in population susceptibility that explains why H5N1 cases are generally detected in younger people than H7N9 (*3,16–18*). We found strong evidence that HA group-matched primary exposure provides profound protection against severe infection and death, for both H5N1 and H7N9, generalizing the concept of ‘original antigenic sin’ (*19*) or ‘antigenic seniority’ (*20*) to address immune imprinting across influenza subtypes. This in turn allowed us to develop new approaches for pandemic risk assessment. Our findings provide new opportunities to advance IAV pandemic preparedness and response, and highlight possible challenges for future vaccination strategies.

### Reconstructing IAV exposure history by birth year

To investigate whether an individual’s initial childhood exposure to IAV may influence later susceptibility to H5 and H7 viruses, we estimated the fraction of each birth-year cohort from 1918 to 2015 with first IAV exposure to H1, H2, or H3 - or the fraction still naïve - for each affected country in our study (China, Egypt, Cambodia, Indonesia, Thailand, Vietnam). We estimated the annual probability of IAV infection in children using published age-seroprevalence data (*21, 22*) and then rescaled this baseline attack rate to account for year-to-year variability in IAV circulation intensity (Supplementary Text).

One resulting country-specific reconstruction of HA history is depicted in Fig. 1B. While H3N2 has dominated since 1968, considerable proportions of many birth-year cohorts from the 1970s onwards were exposed first to H1N1 viruses, with notable peaks near the reemergence of H1N1 in 1977 and the emergence of the 2009 H1N1 pandemic virus.

**Fig. 1.**
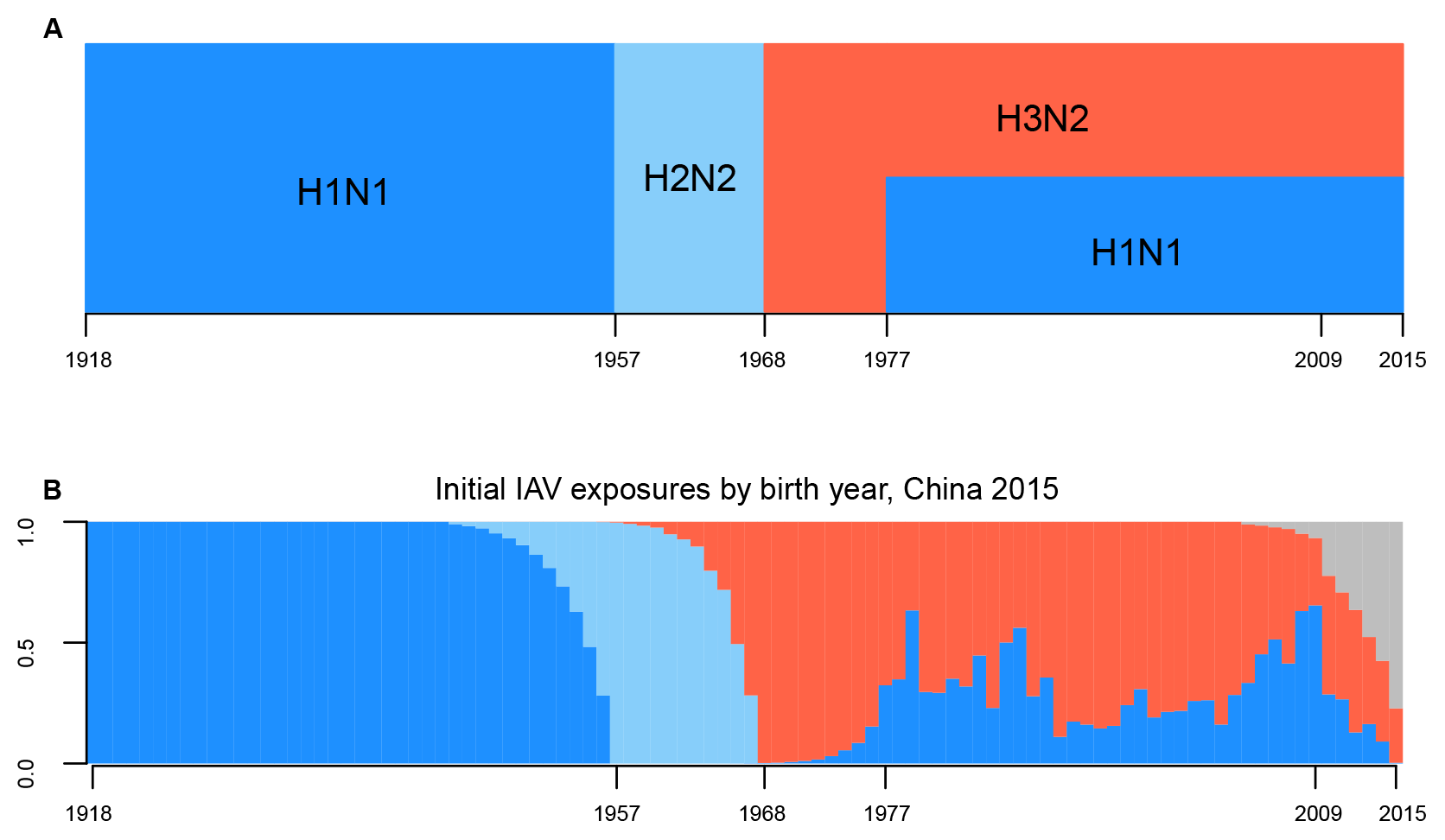
Reconstruction of 20^th^ century HA imprinting. (**A**) History of seasonal IAV circulation, and (**B**) estimated fraction of each birth cohort in China with initial exposure to each subtype, as of 2015. Estimated patterns in other countries (not shown) are identical up to 1977, and very similar thereafter. In both panels, pandemic years are marked on the horizontal axis. Blue represents group 1 HA viruses, red represents group 2, and grey represents naïve children who have not yet experienced an IAV infection.

### H5N1 and H7N9 cases track HA imprinting patterns

Next, we compiled data on all known human cases of H5N1 and H7N9 with reported patient age (Fig. 2A, Fig 2B). These data encompass mostly clinically severe and fatal cases; total incidence remains unknown, particularly for H7N9, where large numbers of undetected, mild cases have been hypothesized (*2*). Thus, our analysis focused on the determinants of severe cases. The possible existence of many mild cases, of any age distribution, is consistent with the HA imprinting hypothesis because partial immunity against influenza is known to reduce the severity of infection without preventing infection altogether (*5,23–26*).

The preponderance of observed H7N9 cases among older cohorts, and H5N1 cases among younger cohorts, is clearly evident (Fig. 2A, Fig 2B). These patterns reflect birth year, not age: H5N1 cases occurred over 18 years from 1997-2015, yet cases from all years exhibit similar dependence on birth year. Analysis of 361 H5N1 cases in Egypt, the one country with a large number of cases across the last decade, shows no trend in case birth years through time, while case age increased steadily (p=0.0003; Fig. S1). So, on average, the same birth cohorts remained at high risk of severe infection, even as members grew ten years older.

Fig. 2C and Fig 2D depict the case data normalized to account for the demographic age distribution in affected countries. If all birth cohorts had equal risk of severe infection, case incidence would be proportional to the demographic age distribution. Bars above the midline thus represent birth years showing excess risk relative to this null expectation, while bars below indicate a shortfall. This correction for demography highlights two points: first, the incidence and mortality data for H5N1 and H7N9 are near-mirror images of each other. Second, the group 1 to group 2 HA transition in 1968 is the key inflection point, with those born before the emergence of H3N2 showing protection against severe cases of H5N1 but not H7N9, and those born after 1968 showing largely the opposite pattern. With H7N9, not only is the inflection point near 1968 clearly evident, but so is the rise in severe cases with birth years near resurgent H1N1 periods around 1977 and 2009. One-sided binomial exact tests showed that excess H5N1 incidence, relative to the demographic null expectation, had a lower probability of occurring in cohorts born before 1968 (p<1e^−10^), while excess H7N9 incidence was more probable in these same cohorts (p<1e^−9^). The same pattern held for excess mortality (Supplementary Text). This suggests that the immune system imprints on conserved HA epitopes from the first-ever exposure to IAV, resulting in heterosubtypic (but within-group) protection against severe infection - akin to antigenic seniority within a single HA subtype (*13,19,20*).

Even more striking is the tight correspondence between the incidence of (and mortality from) H7N9 and H5N1 and the *a priori* prediction based on HA imprinting patterns and demographic age distributions (Fig. 2). We emphasize that the black lines in Fig. 2 are not fitted to the case data, but are independent predictions (Fig. 1B). Differences between the predictions and data are remarkably small, and likely arise from known aspects of influenza epidemiology not incorporated in the predictions (see below), generalization across time and countries (e.g. attack rates for the reconstruction came from German data, but focal populations are largely Asian), and noise arising from small case numbers. In contrast, NA imprinting patterns are a poor fit to H5N1 and H7N9 case data (Fig. S2), and NA-mediated protection is not supported by statistical modeling.

**Fig. 2.**
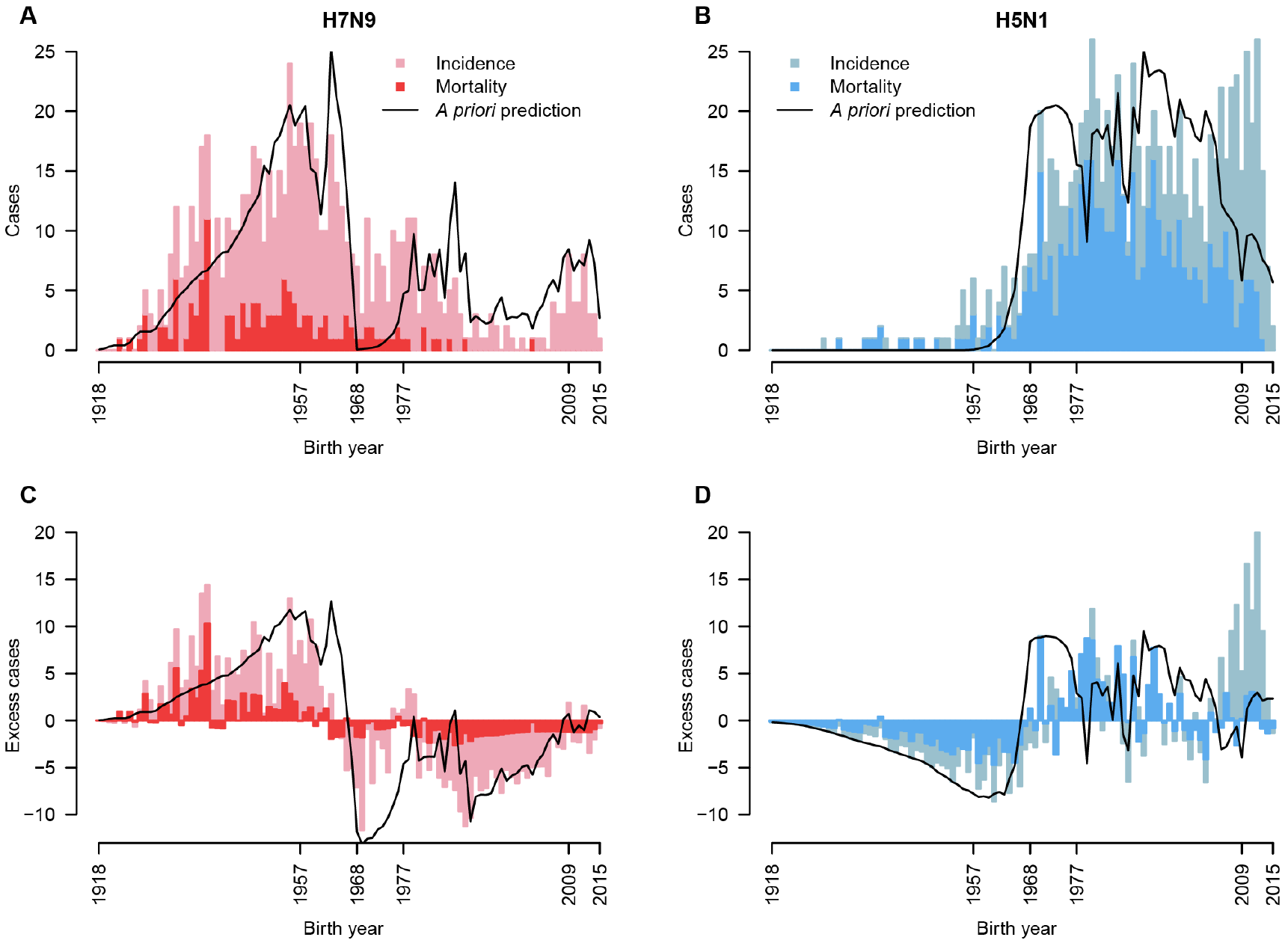
H7N9 and H5N1 observed cases and deaths by birth year. Bars show observed data, black lines show *a priori* prediction based on demographic age distribution and reconstructed patterns of HA imprinting (Methods). (**A**) Birth year distribution of 680 H7N9 cases, from China, 2013-2015, and (**B**) 835 H5N1 cases, from Cambodia, China, Egypt, Indonesia, Thailand and Vietnam, 1997-2015. (**C, D**) Aggregate case data normalized to demographic age distribution from the appropriate countries and case observation years.

### HA imprinting explains age distributions

To formally assess the HA imprinting hypothesis alongside previous explanations (*2,4–8*) for observed H5N1 and H7N9 age distributions, we developed a set of multinomial models. These models related the probability that a case occurred in a given birth cohort to country-and year-specific demography and risk factors including age-based risk of exposure to poultry, age-based risk of severe disease or case ascertainment, and reconstructed patterns of first exposure (and hence potential imprinting) to HA or NA subtypes (Table S1). Model comparison showed that HA imprinting was the dominant explanatory factor for observed incidence and mortality patterns for both H5N1 and H7N9 (Table 1, Figs. S3, Fig S4, Table S2), as it was the only tested factor included in all plausible models for both viruses (i.e. all models with Akaike weights greater than 4e^−5^).

The best models also accounted for age-based risk of severe disease. Age-based poultry exposure risk (estimated based on contact data from China (*7,8*) was included for H7N9 but not H5N1, perhaps reflecting that age-specific poultry exposures vary across the multiple countries affected by H5N1 or that humans interact differently with ill (H5N1-infected) versus asymptomatic (H7N9-infected) poultry. In models including HA imprinting, NA imprinting never showed any significant effect (Table S2). Remarkably, despite differences between the viruses and age cohorts involved, the estimated protective effects of HA imprinting were nearly identical for H7N9 and H5N1. In all models, protective HA imprinting reduced the risk of severe infection with H5N1 or H7N9 by ~75%, and the risk of death by ~80% (Table 1, Figs. S3–Fig S5, Table S2).

**Table 1.** Estimated protection from HA imprinting. D = Demography, E = Exposure to poultry, A = High-risk age groups, H = HA imprinting, N = NA imprinting (see Methods, Table S1).

| Factors in model* | HA imprinting protection (95% CI) | $\Delta AIC$ | Akaike weight |
| --- | --- | --- | --- |
| **H5N1** |  |  |  |
| DAH | 0.75 (0.65-0.82) | 0.00 | 0.9994 |
| DEAH | 0.83 (0.76-0.88) | 15.35 | 4.65E-4 |
| DEANH | 0.83 (0.73-0.88) | 17.32 | 1.74E-4 |
| DH | 0.80 (0.71-0.85) | 69.18 | 9.50E-16 |
| DEH | 0.87 (0.80-0.90) | 103.31 | 3.69E-23 |
| DENH | 0.86 (0.78-0.90) | 105.29 | 1.37E-23 |
| **H7N9** |  |  |  |
| DEAH | 0.76 (0.67-0.82) | 0.00 | 1.00 |
| DAH | 0.81 (0.74-0.87) | 42.87 | 4.09E-10 |
| DEH | 0.84 (0.78-0.88) | 61.59 | 4.23E-14 |
| DENH | 0.83 (0.75-0.88) | 62.26 | 3.02E-14 |
| DH | 0.88 (0.84-0.92) | 138.40 | 8.83E-31 |

### Antigenic seniority across influenza subtypes

Most individuals born before the emergence of H3N2 in 1968 have been exposed to H3N2 after 1968, probably multiple times, yet these seasonal group 2 exposures later in life evidently fail to override group 1 HA imprinting from childhood (Fig. 2). This pattern indicates that a form of antigenic seniority can act across HA groups (not just within subtypes, as conventionally assumed). It will therefore be desirable to determine precisely how first HA exposure trumps subsequent HA exposures in eliciting protection (i.e. the precise mechanism underlying antigenic seniority in this context). One intriguing possibility was highlighted by the 2009 H1N1 pandemic, which demonstrated the importance of a heterosubtypic immune mechanism consistent with protection at the level of HA groups: IAVs containing antigenically unfamiliar HA head domains, but familiar (based on previous exposure to an HA in the same group) conserved HA stem domains (*9–15*), can preferentially boost broadly cross-protective antibodies against the conserved domains. Specifically, exposure to H5N1 or H7N9 can activate HA stem-specific reactivities induced by previous infection by H1 or H3, respectively (*14,15,27*). Together with these observations, our results support antigenic seniority acting on B-cell-mediated immune responses targeting conserved HA epitopes (shared within but not between HA groups) as a plausible mechanism underlying the birth-year specific protection seen for human H5N1 and H7N9 infections. NA imprinting clearly does not explain these patterns (Fig. S2, Table S2), and other potential antigens (e.g. the nucleoprotein or the extracellular domain of the matrix 2 protein) have not experienced pandemic reassortment events since 1918.

Protein sequence and structure data are consistent with a stem-directed mechanism for antigenic seniority acting at the HA-group level (*28*). Amino acid sequences in the HA stem exhibit divergence within a phylogenetic group that is comparable to divergence in globular head sequences within a single HA subtype like H1 (i.e. the scale at which antigenic seniority is already known to act (*20*); Fig. S6). Given that H3 and H7 are as divergent as any pair of group 2 HAs, H3 childhood exposure may protect as well or better against the other group 2 HAs (H4, H10, H14, H15), but perhaps not at all against the considerably more divergent group 1 HAs (Fig. S6C). Similarly, H1 or H2 childhood exposure may protect generally against most or all group 1 HAs but not against group 2 HAs. However, our empirical findings are also consistent with cross-reactivity to conserved HA head epitopes shared within, but not between, HA groups (*29*). Given the immunodominant nature of HA head reactivities (*10,15,30*), the possible involvement of conserved HA head epitopes (or both head and stem epitopes) should be evaluated. Cross-reactive HA-specific CD4+ or CD8+ T cell responses should also be investigated, since they are also likely to be disproportionately shared within HA groups (given the sequence similarities within each clade) and might be especially capable of facilitating long-term protection as indicated by the data.

### Rational projections of future pandemic risk

For any new pandemic IAV strain (i.e. one that has acquired a capability for efficient human-to-human transmission), epidemiological impacts of the first pandemic wave would be shaped not only by age-based mixing patterns (*31–33*) but also by HA imprinting patterns from past IAV circulation. We created projections for a putative pandemic-capable H5 or H7 strain - such as a gain-of-function strain or a natural variant with mutations increasing human-to-human transmissibility - possessing the same antigenic properties as the current H5 or H7 strains. The data on observed H7N9 and H5N1 cases enabled us to quantify how matched HA imprinting reduces the probability of developing a severe infection, but not how matched imprinting affects an individual’s probability of acquiring a milder infection or the infectivity of such mild infections. People who become infected despite prior immunity likely transmit influenza at reduced rates owing to diminished viral titers and viral shedding, as observed in humans and animal models (*5,23–26*). We thus made the conservative assumption consistent with these findings that imprinting does not change the probability of acquiring infection upon exposure, but can reduce severity and infectivity in individuals with protective HA imprinting.

Fig. 3A illustrates the projected age-structured attack rate of severe cases for hypothetical pandemics of H5 or H7 IAV occurring in 2015 in the United Kingdom. The projected risk profiles for severe infection are shaped strongly by HA imprinting, including the prediction that individuals above 50 years of age (i.e. born well before 1968 and first exposed to a group 1 HA) would experience much lower morbidity than younger age groups in an H5 pandemic. Similar projections for China and Vietnam reveal the influence of demographic differences between countries (Fig. S7). The qualitative patterns in projected age impacts are robust to the assumed infectivity of mild cases arising in individuals with protective HA imprinting (Fig. S7A).

Projections for pandemics occurring a decade from now highlight predictable shifts in severe disease risk patterns as the imprinted population ages, with the key pivot point around birth years near 1968 shifted to older ages (Fig. S7). Impacts in the youngest age groups, on the other hand, depend on patterns of IAV circulation in the coming decade (Fig. S7). All pandemic projections that account for HA imprinting exhibit markedly lower severe attack rates than projections that assume no protection from imprinting (Fig. 3A, Fig. S7). Total attack rates (including mild and severe cases) are higher and more evenly distributed across age groups than the severe attack rates shown here.

**Fig. 3.**
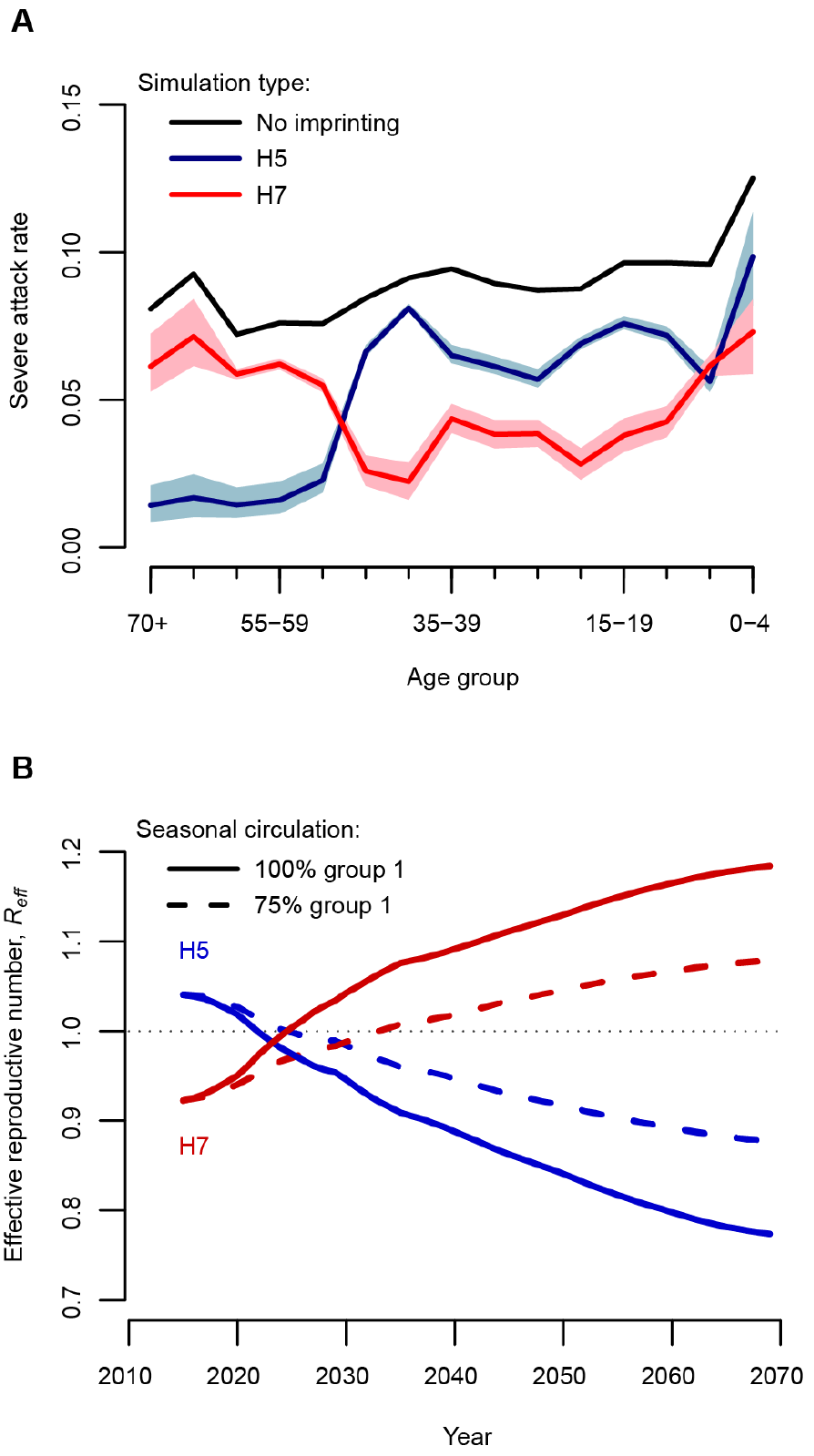
Projected effects of HA imprinting on future pandemics. (**A**) Attack rate of severe cases, by age group, for hypothetical H5 (blue) and H7 (red) IAV pandemics in the UK in 2015 (*R*_0_=2.5, relative infectiousness of imprinting-protected individuals (*α*) = 0.5). Simulations assume UK demography and age-structured mixing, HA imprinting effects, and elevated risk of severe infection in young children and the elderly (Supplementary Text). Colored lines show the average outcome, and shaded regions include 95% of 100 simulated outcomes. Black lines show outcomes assuming no HA imprinting. (**B**) Projected change in *R*_*eff*_, for hypothetical H5 (blue) or H7 (red) IAV with *R*_0_=1 2 and *α*=0.5, if group 1 IAVs make up 100% or 75% of seasonal circulation after 2015.

Over any prolonged period when IAV circulation is dominated by one HA group, imprinting generates growing herd immunity against zoonotic IAV strains from that group. Conversely, zoonotic strains from the mismatched HA group benefit from the rising proportion of humans without protection. So long as mild cases arising in people with group-matched imprinting contribute any less to transmission than unprotected cases, or if some fraction of infection events are prevented by imprinting-derived immunity, then imprinting will inevitably alter the transmissibility of zoonotic IAV strains in the human population. This is summarized by the effective reproductive number, *R*_*eff*_, which quantifies transmission in a partially immune population (Fig. 3B). Crucially, a zoonotic strain that is initially subcritical (with *R*_*eff*_ < 1 and therefore unable to spread sustainably) can - due solely to susceptibility changes in the human population - emerge as supercritical, and hence as a pandemic threat, if the mismatched HA group dominates IAV circulation for a sufficient period (Fig. 3B).

Interestingly, the co-circulation of group 1 and 2 HAs since 1977 has balanced herd immunity in a way that could reduce the probability that supercritical zoonotic strains from either HA group emerge (i.e. by preventing an otherwise larger buildup of those susceptible to one or the other group). As a generality, *R_eff_* for zoonotic influenza strains will change through time depending on seasonal influenza patterns and demographic background, and the magnitude of change will depend on the infectivity of imprinting-protected cases (Fig. S8).

## Discussion

Our findings show that major patterns in zoonotic IAV epidemiology, previously attributed to patient age, are in fact driven by birth year. IAV strains circulating during an individual’s first years of life confer pronounced, long-term protection against novel HA subtypes from the same phylogenetic group. Hence, the phenomenon of antigenic seniority extends across IAV subtypes, introducing previously unrecognized generational structure to influenza epidemiology. Influenza virulence represents a joint phenotype between virus and host - even for strains not yet adapted to the human population - and includes a major contribution from immune imprinting. This has several implications for public health and raises some interesting questions.

These findings support the hypothesis that the unusual mortality pattern during the 1918 influenza pandemic - when young adults who had imprinted on an H3 (group 2) HA in childhood suffered severe outcomes from an H1 (group 1) IAV - may have been due primarily to HA group-mismatched first exposure in this cohort (*18*). This same cohort (born before ~1900) was strongly affected during the mismatched H2 (group 1) 1957 pandemic (*34*); yet they suffered no excess mortality when they were even older, during the 1968 H3 (group 2) pandemic (*35*). We speculate this effect may extend to seasonal H1N1 and H3N2 viruses (i.e. that H3N2’s current greater health impact is driven at least partially by today’s older age classes, already intrinsically at higher risk for IAV morbidity, having imprinted on a group-mismatched HA). Hence, a diagnostic assay able to determine whether a person’s first IAV infection involved a group 1 or group 2 HA may be useful for individualized clinical care and even vaccine design strategies, both for pandemic and seasonal IAVs: in other words, our results strengthen earlier proposals (*36*) that influenza vaccines tailored to specific exposure histories is an idea worth exploring.

We cannot yet be certain whether similar imprinting protection (and predictive power) extends across all the potential emerging HA subtypes, as sufficient epidemiological data exist only for H5N1 and H7N9. Yet the fact that H3 and H7 are as distantly related as any pair of group 2 HAs can be (Fig. S6), and are still involved in cross-protection, suggests HA imprinting effects would likely influence other subtypes. This position is supported by the preponderance of mismatched childhood HA exposures among the handful of clinically significant human cases of other novel subtypes detected to date: pooling data from 28 human cases of H5N6, H6N1, H7N7, H9N2, H10N7 and H10N8, age patterns are consistent with HA imprinting (p=0.019; see Supplementary Text), but case numbers are insufficient to investigate particular subtypes. Immunological experiments (e.g. using chimeric HA proteins (*14*)) are needed to systematically map HA cross-protection patterns across all HA subtypes, both within and between HA groups.

Our findings also raise questions about whether seasonal influenza vaccination might boost broadly-protective anti-HA responses or otherwise alter imprinting from natural infection. The persistence of group 1 imprinting in older adults despite decades of natural exposure to H3N2 after 1968 (Fig. 2), and the relative weakness of group 2 anti-HA stem reactivities in these older cohorts (*13*), suggest that HA exposures later in life do not readily alter the sort of strong protection we detect in those already imprinted to a particular HA group. However, it is plausible that vaccination of very young, IAV-naïve children could interfere with imprinting. Specifically, by exposing IAV-nai’ve children simultaneously to group 1 (H1N1) and group 2 (H3N2) antigens, vaccination might prevent strong imprinting against either HA group, or confer dual imprinting to both HA groups - or it could have no effect beyond delaying the age of imprinting via the first natural infection. Importantly, sensitivity analyses demonstrated that, given the low IAV vaccination coverage in H5N1-and H7N9-affected countries, none of these possible effects would appreciably change the conclusions of our study (Fig. S5). However, to properly inform early childhood vaccine policy, future work is necessary to determine which, if any, of these effects occur.

Another obvious implication of HA group imprinting is that it might complicate ‘universal’ vaccination approaches that target conserved HA stem domain epitopes. Our findings indicate potent, long-lasting cross-protection between subtypes, putatively based on such responses. However, to generate protective anti-HA stem responses to both HA groups, universal vaccination may have to outperform natural infection in its ability to induce broad immunity in the face of previous imprinting given that H3N2 exposure fails to produce discernible, long-lasting immunity in individuals who first encountered group 1 HA viruses. To be effective, would bivalent (group 1 and group 2 HA stem) universal vaccines need to be delivered to infants prior to natural IAV infection? Or, might universal vaccines impair natural, long-term protection of the sort we have detected against H5N1/H7N9 if received prior to an individual’s first natural IAV infection?

Our work implies that we have never seen a true ‘virgin soil’ influenza pandemic, and therefore all prior estimates of the basic reproductive number, *R*_0_, for pandemic IAVs are systematic underestimates since they do not account for protection induced by HA imprinting. Projections of age-impact patterns and *R*_*eff*_, like the ones we present here, could be done systematically for any country with contact and demographic data. This would provide rolling estimates of which age groups would be at highest risk for severe disease should particular novel HA subtypes threaten to emerge. These projections could help to guide cohort-targeted or region-specific prevention, preparation, or control efforts, and can be built from present knowledge. To achieve quantitative projections of changes in *R*_*eff*_, we will need to determine the epidemiological nature of protection arising from matched imprinting: is some fraction of cases prevented entirely, and by what factor is infectivity reduced in mild cases arising in protected individuals? Notably, there are indications that H5N1 may infect individuals with matched imprinting considerably less frequently than does H7N9 (*2*). Intensive household investigations, including cases and contacts with matched and mismatched imprinting, and serological screening for subclinical infections, will yield crucial information.

More generally, our findings show that emergence risk cannot be considered in isolation, even for ‘novel’ pathogens that have not circulated in humans before. These pathogens are commonly assumed to have a blank slate of immunologically naïve humans to infect, but cross-protection arising from exposure to related pathogens can generate substantial levels of population immunity, such that the emergence process is governed by bottom-up control. When the community of related pathogens undergoes changes, as when influenza pandemics cause shifts in seasonal strain dominance, the landscape of population immunity will change accordingly. For influenza A viruses, population immunity is evidently a major determinant of observed epidemiological patterns, and ultimately of the risk of pandemic emergence. Because population immunity changes predictably, a major component of emergence risk is thus quantitatively predictable.

## Methods

### Case data

We compiled data describing all reported human cases of H7N9 (2013-Nov. 2015) and H5N1 (1997-Nov. 2015) influenza. We obtained case data from three previously published H5N1 line lists, one spanning 1997 (*37*) and the other spanning Sept. 2006-Aug. 2010 (*38*), as well as one H7N9 line list spanning Jan.-Sept. 2013 (*39*). For all other cases, we compiled original line lists using reports from the WHO as our primary resource and the Hong Kong Centre for Health Protection as a secondary resource. We incorporated additional cases or case details from Flu Trackers. Our full database included 842 lab-confirmed cases of H5N1 between 2003 and Nov. 2015, and 545 lab-confirmed cases of H7N9 between Oct. 7, 2013 and Nov. 2015. These closely match numbers of confirmed cases reported by the WHO over the same time period: 844 cases of H5N1 and 546 cases of H7N9. Our line lists are available as supplementary data files. Hyperlinks to each case information source are provided within.

We included confirmed, suspected and probable cases in our final analysis, but verified that our results are robust to the exclusion of unconfirmed cases (Fig. S5). We excluded cases for which patient age was not reported. To date, all reported human cases of H7N9 have occurred in China. Of H5N1 cases with patient age reported, 95% occurred in Cambodia, China, Egypt, Indonesia, Thailand, and Vietnam, so we analyzed only cases occurring in these countries. In total we analyzed 835 H5N1 cases and 440 deaths, and 680 H7N9 cases and 132 deaths. Mild or asymptomatic cases of both infections may not be ascertained (*2*), so our results should be interpreted only as predictors of severe, symptomatic infection.

### Normalization of data to demographic age distribution

If all birth years were at equal risk of severe infection, we would expect the observed age distribution of cases to be proportional to the demographic age distribution. To examine excess incidence of severe infection or death, relative to this demographic null expectation, we normalized the observed data to the demographic age distribution (Fig. 2C, Fig 2D). We tabulated the observed and expected number of cases occurring in each birth year, *i*, for each country, *c*, in each case observation year, *y*. To determine birth year, we subtracted case age from the year in which case onset occurred. To estimate the expected number of cases in each birth year, *Exp*_*yci*_ we used the following formula:

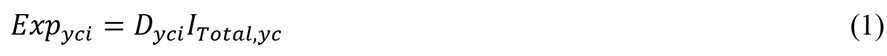

Here *D*_*yci*_ describes the fraction of the total population, in country *c*, year *y*, that belongs to birth year *i*. *lTotai,_yc_* describes the total number of cases or fatalities that occurred in yeary, country *c*.

Finally, we defined normalized values within each birth year, *N*_*i*_ as the unweighted sum, across all possible country-y I’m in my lab ears, of differences between observed and expected numbers of cases or fatalities:

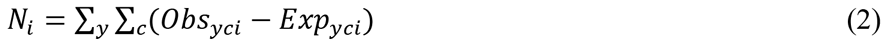

### Model formulation

We used multinomial models to describe the probability distribution of H5N1 or H7N9 cases or fatalities across birth years. The multinomial distribution requires a set of parameters, *p*_*yci*_, which described the probability that an infection observed in year *y*, country c, occurred in birth cohort i. Within each model, a unique combination of factors, including age-based risk of severe infection, poultry exposure risk, HA imprinting, and NA imprinting, determined *p*_*yci*_. For each virus (H5N1 and H7N9), we fitted models independently for each case outcome (infection or death).

For each candidate model, we computed maximum likelihood parameter estimates to quantify the effects of relevant explanatory factors on birth-cohort risk (Table S1). Parameters *H_m_* and *N_m_* described the relative risk of infection or mortality for those with protective HA or NA imprinting, while parameters *A_c_* and *A_e_* described relative risk for young children (<5 years old) and the elderly (>65 years old).

We performed model comparison using AIC to determine which combination of factors best explained observed distributions of H5N1 or H7N9 incidence or mortality across birth years. We calculated Akaike weights, *w_z_*, which can be interpreted as the proportional evidence in support of model *z* as the best of all models tested. Weights are calculated using the expression, 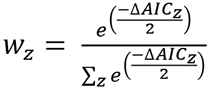(*40*). In many cases, when we added NA imprinting to models already considering HA imprinting, the maximum likelihood estimate of the *N_m_* parameter was 1. In these cases, NA imprinting had no effect, and the new, more complex model became identical to the simpler model. We excluded these degenerate models from Akaike weight calculations.

### Factors tested

#### Demography (D)

If all individuals were at equal risk of infection, regardless of birth year, the number of cases occurring in a given birth year would be proportional to the fraction of the population born in that year, as given by the country’s demographic age distribution. Thus, for every country, *c*, and case observation year, *y*, a vector, *D*_*yc*_, described the demographic age distribution and served as the null predictor of infection risk in each birth cohort. Vector *D*_*yc*_ was normalized to sum to 1, so that each element described the fraction of the population born in a particular year. Using the US Census Bureau’s International Database (*41*), we obtained publicly available demographic data for each country and year of case observation. Demography was included as a factor in every model tested (Supplementary Text).

#### Exposure to poultry (E)

Though limited human-to-human transmission is thought to occur, the majority of H7N9 and H5N1 infections are caused by spillover from infected poultry (*2, 3*). To build models incorporating age-based patterns of exposure to poultry (E), we obtained published data from three urban locations (Guangzhou, Shanghai and Shenzhen), and two non-urban locations (subrural Guangzhou and Xiuning) in China (*7, 8*). Note that we included data from two independent surveys conducted in urban Guangzhou, for a total of six survey data sets. No poultry exposure data were available for other affected countries.

Although poultry exposure patterns vary by location, even within China, large reported uncertainties and geographic constraints on the available data prohibited a statistically valid effort to match cases with geographically specific exposure patterns. Instead we computed an average exposure rate for each birth cohort and year of H5N1 or H7N9 case observation, as follows. Each survey reported poultry exposure rates by age group. We used these values to assign each birth cohort an age-specific poultry exposure rate, based on their ages in each possible year of case observation. If exposure rates were not reported explicitly for young children, we substituted the rate reported for the youngest available age group. We then normalized across birth cohorts to determine the proportional risk of exposure at each survey location. Normalization ensured that survey locations reporting higher overall rates of exposure did not disproportionately influence model inputs. Finally, for each birth year, we took the average proportional exposure rate across all six poultry exposure surveys.

#### Age-based risk of severe infection (A)

A basic principle of flu epidemiology is that children under 5 and elderly adults over 65 are high-risk groups for influenza infection (*42–44*) These groups may be more susceptible to severe influenza infections, or may seek healthcare at an increased rate, leading to case-ascertainment biases. Models including the age-based risk factor (A) introduced two parameters that allowed each high-risk age group to experience increased risk relative to adults and children ages 5-64. Parameter *A*_*c*_ quantified the relative risk of young children, while parameter *A*_*e*_ quantified the relative risk of the elderly (Table S1). Both parameters are constrained take a minimum value of 1 so as to represent elevated risk of severe infection.

For mortality analyses, we noted that case-fatality rates are not always elevated in children, despite their elevated risk of severe infection (*43*). Thus, for mortality analyses we relaxed the constraint on parameter *A*_*c*_, allowing the data to inform whether young children exhibit increased (*A*_*c*_ > 1) or decreased (*A*_*c*_ < 1) risk of death from H7N9 or H5N1 infections, relative to the general population.

#### Hemagglutinin imprinting (H)

Models that considered hemagglutinin imprinting (H) divided the population into two exposure groups: those with group-matched first IAV exposures (protective HA imprinting), and all others (non-protective HA imprinting or naïve children). We calculated the fraction of each birth cohort with first exposure to either HA group using methods described below (see Reconstructing immune imprinting patterns). We initially separated naïve children from adults with non-protective HA imprinting, but our results consistently indicated no difference in risk between these groups (results not shown), leading us to combine naïve children and mismatched HA imprinting into a single reference group.

Hemagglutinin imprinting models introduced a single parameter, *H*_*m*_, which allowed the group with protective HA imprinting to experience decreased risk relative to all others in the population.

#### Neuraminidase imprinting (N)

As for HA imprinting, models that considered NA imprinting divided the population into two groups: one with matched (protective) first IAV exposures, and a reference group that included all others. Models containing factor N introduced one parameter, *N*_*m*_, which allowed the group with protective NA imprinting to experience decreased risk. Two phylogenetic groups have been identified for NA, as for HA. We are not aware of experimental evidence of cross-protection between NA subtypes within the same phylogenetic group (without which NA history could not explain incidence patterns for H7N9, since N9 is not known to have circulated in the human population). However, given that N9 and N2 fall in the same NA group, we considered the possibility that imprinting on N2 might be protective against N9. H5N1 is clearly matched to seasonal H1N1, as they share the N1 subtype.

### Models tested

We tested models that considered all possible combinations of the above four factors, as well as demographic age structure. Our full set of 16 models included: D, DE, DA, DH, DN, DNH, DAH, DAN, DANH, DEA, DEH, DEN, DEAH, DEAN, DENH, and DEANH. For each model, we fit relevant parameters to the appropriate data before performing model comparison. We repeated each model analysis using both incidence and mortality data for H7N9 and H5N1. See Supplementary Text for model equations and likelihood functions.

### Reconstructing immune imprinting patterns

To inform hemagglutinin and neuraminidase imprinting models, we estimated the fraction of each birth cohort with first IAV exposure to seasonal subtypes H1N1, H2N2 or H3N2, and the fraction that remained naïve.

We first used a truncated geometric model to estimate the baseline probability that first IAV infection occurs at a given age. Because age-seroprevalence studies report that 98-100% of children have been infected with IAV by age 12 (*21, 22, 45*), we set 12 as the maximum possible age of first infection. Let *ε_ij_* be the probability that an individual with birth year *i* has his or her first IAV infection in calendar year *j*. Then:

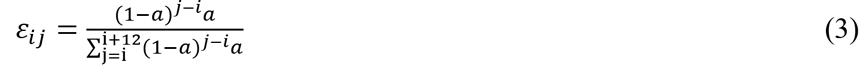

where *a* is the annual probability of infection (the baseline annual attack rate on seronegative children), and *j* takes values from *i* to *i*+*12*. Using published age-seroprevalence data (*21, 22*), the maximum likelihood estimate for *a* was 0.28 (95% CI 0.26-0.30), consistent with other IAV attack rates estimated in children (*45–47*).

To account for variability in the annual attack rate between years, we compiled an index of IAV circulation intensity from 1918-2015 (Supplementary Text). We defined the scaled annual attack rate as *a*_*m*_ = *aI*_*m*_, where *m* is the calendar year, *I*_*m*_represents that year’s intensity score and *a* represents the baseline annual attack rate estimated above.

Finally, we modified equation 3 to account for variable annual attack rates and the possibility that some children have not yet been exposed:

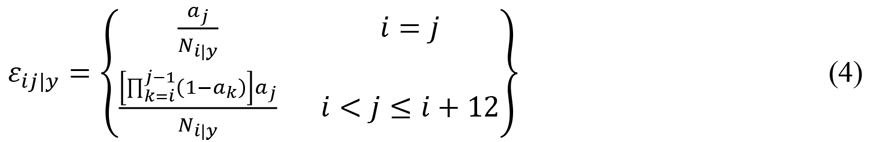

Here, *ε_ij|y_* is the probability that individuals in birth cohort *i* experienced their first infection in calendar year *j*, given that H5N1 or H7N9 case observation occurs in year *y*. Note that because birth cohorts are very large, the fraction of birth cohort *i* with first exposure in year *j* converges to probability *ε_ij|y_*. The expression, 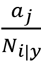 represents the probability of infection in the first year of life (at age 0), and the expression 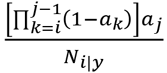 represents the probability of infection at ages 1-12. The normalizing factor, *N*_*i|y*_, reflects the assumption that all individuals have had their first infection by age 12, and is used to ensure that all relevant probabilities for an age group sum to 1. It is taken as:

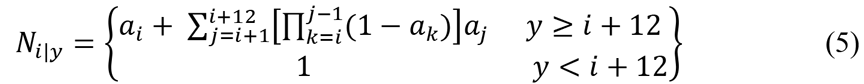

When the birth cohort is older than 12 in the year of observation, *N*_*i|y*_ always falls within the range (0.98-1.00) and minimally affects the final outcome. For birth cohorts that are younger than 12 at the year of observation (*y*<*i*+12), normalization is not appropriate because some individuals in the age group have not had their first exposure; in these instances we set *N*_*i|y*_ = 1 and calculate the naïve fraction as the complement of the cumulative probability of first exposure (equation 7).

Finally, we combined the age of first infection probabilities with seasonal circulation patterns to determine the fraction of age group *i* with first exposure to subtype *S* in year *j* and country *c* (*w*_*S, i|y, c*_). We scaled *ε*_*i, j|y, c*_ by the fraction *f*_*S, |j, c*_ of circulating IAVs belonging to subtype *S* (where *S* could represent H1N1, H3N2 or H2N2):

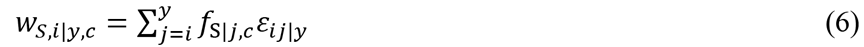

For birth cohorts younger than 12 in the year of case observation (*y*<*i*+*12*), a portion of the cohort had not yet experienced a first IAV exposure. The fraction of cohort *i* that remained naïve in year *y* was given by:

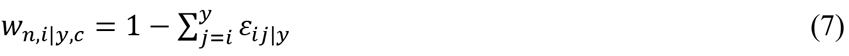

Values of *f*_*S, |j, c*_ were determined by IAV circulation history, and for all *j*, *f*_*H1N1|j, c*_ + *f*_*H2N2|j, c*_ + *f*_*H3N3|j, c*_ = 1 Prior to 1977, a single IAV subtype circulated each year, and pandemic years marked circulation of a new seasonal subtype. Thus, in years 1918-1976, *f*_*s|j,c*_ was set equal to 1 for the circulating subtype, and 0 for all other subtypes.

For years 1977-2015, when H1N1 and H3N2 have circulated simultaneously, we estimated *f*_*S|j, c*_ using influenza surveillance data reported by WHO collaborating laboratories (*48, 49*). For each year of interest, we defined relative incidence as follows: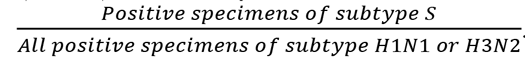. Whenever possible (1997-2015), we used surveillance data from the country of interest (China, Egypt, Cambodia, Indonesia, Thailand or Vietnam) to estimate country-specific relative incidence. When fewer than 50 specimens were reported for a given country in a given year, we substituted surveillance data from the same year, across all other countries of interest with adequate data. For years in which no surveillance data was available from any H5N1 or H7N9 affected country (1977-1996), we substituted surveillance data from laboratories in the United States (*49*). From 1997-2015, the relative incidence of H3N2 or H1N1 in the United States was significantly correlated with subtype-specific relative incidence in our study’s six countries of interest (Pearson’s r = 0.70, p < 0.001), suggesting that US data is an appropriate proxy.

### *A priori* predictions

The *a priori* predictions in Figure 2 illustrate how the observed total number of cases would be distributed across birth years if individuals with protective HA imprinting never experienced severe infection, but risk is otherwise identical for everyone. Thus, the distribution of observed cases across birth years was expected to be proportional to the distribution of unprotected individuals across birth years (i.e. to the demographic age distribution, after individuals with protective HA imprinting had been removed). For each country and year in which cases were observed, this *a priori* prediction is:

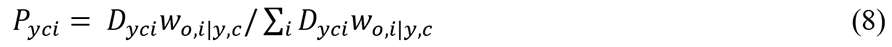

Here, *P*_*yci*_ is the fraction of all cases observed in country *c*, year *y* predicted to occur in birth year *i*. *D*_*yci*_ represents the demographic age distribution as defined in equation 1. Building from equations 6 and equation 7, *w*_*o,i|y,c*_ represents the fraction of each birth cohort with first exposure to a subtype in the opposite HA group as the challenge strain, or naïve to IAV. For the predictions shown in Figures 2A and Figure 2B, we scaled *P*_*yci*_ by the number of cases, *l*_*Total,y,c*_ observed in year *y*, country *c*, and then summed across all affected countries and years: 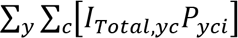. For the predictions shown in Figures 2C and Figure 2D, we subtracted the demographic null expectation (as in equations 1 and equation 2), to obtain the prediction: 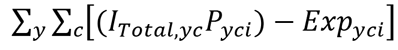.

### Sensitivity analyses

We verified that our conclusions are robust to model assumptions by repeating model fitting and model comparison on eleven variations of the main model described above (Fig. S5). First, to test for robustness against uncertainty on annual intensity of seasonal IAV circulation, we repeated all model analyses assuming a constant annual attack rate (*a*_*j*_ = *a* for all years *j*). Second and third, we considered uncertainty on the baseline attack rate, *a*, by setting this parameter equal to its upper and lower 95% confidence bounds. Fourth, we checked for robustness to the estimated annual dominance of H1N1 and H3N2 by fixing the relative incidence of each to its average from 1977 to 2015 (0.34 for H1N1, 0.66 for H3N2). Fifth and sixth, to generate upper and lower bound estimates of either subtype’s seasonal dominance, we increased the observed relative incidence of H1N1 or H3N2 by 0.05 for all years from 1977-2015, while simultaneously decreasing the relative incidence of the complementary subtype by the same amount. Seventh and eighth, we considered two alternate poultry-risk distributions, using data exclusively from Urban Shenzhen (*7*) and Shanghai (*8*) as data from these surveys respectively reported the highest and lowest poultry exposure rates in cohorts born before 1968. Ninth through eleventh, although overall influenza vaccine coverage is low in all six countries considered (approaching 10% in Thailand and <5% elsewhere) (*50*), we tested whether our analysis is robust to childhood vaccination effects. We considered three scenarios, in which vaccination of naïve children could prevent imprinting and leave the child fully susceptible to both HA groups, replace imprinting and leave the child protected against both HA groups or delay imprinting via delay of the first natural infection. To be conservative, these models considered an upper-bound case for IAV vaccination coverage, and assumed childhood coverage levels beyond what is actually achieved in H5N1-and H7N9-affected countries (Supplementary Text). Finally, to verify that our results were robust to the exclusion of data from probable or suspected cases, we repeated model analyses using only laboratory-confirmed cases.

### Phylogenetic and amino acid sequence analyses

We aligned amino acid sequences of representative H1 strains (HA globular head) and group 1 and group 2 HA strains (stem domain) in Geneious v9.0.4 (*51*) (global alignment with free end gaps, BLOSUM62 cost matrix). Maximum likelihood phylogenies were estimated in RAxML version 7.2.8 (*52*) using the aligned amino acid sequences and a GAMMA LG protein model (rapid hill-climbing algorithm). We generated a heat map of HA stem amino acid sequences with percent similarities (BLOSUM62 matrix with threshold 0) calculated in Geneious version 9.0.4 (*51*).

### Projections of future pandemics

We used a discrete-time stochastic model to create projections of the age-specific severe attack rates in a hypothetical future H7 or H5 pandemic. We conducted simulations for the United Kingdom, China, and Vietnam in 2015 and 2025. For the 2025 simulations, we considered two scenarios for seasonal IAV circulation between 2015 and 2025: 25% group 1 and 75% group 1 circulation. Demography and imprinting patterns for each country and year were obtained as described in Methods (see ‘Normalization to demographic data’ and ‘Reconstructing immune imprinting patterns’).

In the model, individuals with matched imprinting have probability (1 - *H*_*m*_) of experiencing a ‘protected’ course of infection and a probability *H*_*m*_ of experiencing the same ‘unprotected’ infection as individuals with mismatched imprinting. ‘Protected’ individuals have some degree of acquired immune protection and do not experience severe disease; to be conservative, we assume that they still become infected but their lower viral loads lead to reduced infectiousness relative to their ‘unprotected’ counterparts (mediated by a relative infectiousness parameter, *α*). As discussed in the main text, this assumption mirrors data from experimental infections in humans and non-human animals, which show that infections in partially-protected individuals exhibit lower viral titers, lower and more short-lived viral shedding, and lower rates of transmission than infections in naïve individuals (*23, 25, 26*). In our analyses, we tested values of *α* spanning from 0.1 to 0.9, representing the full range of possible reductions in transmission. We expect that the true value for a given situation would depend on specific properties of the focal IAV strain as well as the past strains that generated the imprinted responses.

An ‘unprotected’ infected individual in age group *i* exposes a total of *X*_*ij*_ individuals from age group *j* to the virus, where *X*_*ij*_ follows a negative binomial distribution with dispersion parameter *k* = 0.94 (*53*) and mean *M*_*ij*_. For ‘protected’ infected individuals, the mean number of new infections is instead *αM*_*ij*_. The matrix **M** (with entries *M*_*ij*_) is the next generation matrix with dominant eigenvalue equal to the pathogen’s basic reproductive number (*R*_0_). *R*_0_ was set at 2.5 so that the effective reproductive number (*R*_*eff*_), which accounts for protection from matched imprinting, was approximately 1.9 at the beginning of the 2015 simulations. This value aligns with the *R*_*eff*_ calculated at the beginning of previous pandemics (*54–58*).

An unscaled next generation matrix **M** was constructed from data on contact rates between age groups. Separate contact matrices giving the relative rates of contacts between individuals in different age groups were used for each country. For the UK, we used the matrix of all reported physical and non-physical contacts in Great Britain reported by Mossong et al. (*31*); for China, we used data on individual-level contact data reported by Read et al. (*32*); and for Vietnam, we used contact diary data from Horby et al. (individual-level data provided by Peter Horby) (*33*). We converted the China and Vietnam contact data from the reported age bins into the age bins needed for simulations by disaggregating the reported data (scaled by demography) into separate age years and then reaggregating the data into the appropriate age bins. All elements of this matrix were rescaled by a constant factor so that its dominant eigenvalue was equal to the desired *R*_0_.

The *X*_*ij*_ exposures caused by an infectious individual were assumed to occur after a serial interval of *T* days from the source case, where *T* was distributed according to a Weibull distribution with mean 3.6 (shape = 2.3 and scale = 4.1) (*59*). Given exposure on day t, the probability that an individual in age group *j* developed the disease was equal to the proportion of age group *j* susceptible on day *t*.

Each individual that experienced an ‘unprotected’ course of infection had a chance of developing a severe infection. The probability of severe infection is a property of a given viral strain (roughly, its virulence) and will vary among scenarios; we chose a baseline value of 0. 1, and note that different choices for this value simply rescale the results and do not alter any qualitative patterns. To account for the elevated risk of severe infection in children (0-4 years) and the elderly (65+), the probability of severe infection in these age groups was multiplied by the age-specific risk parameters *A*_*c*_ and *A*_*e*_. We used consensus parameter estimates for *A*_*c*_ and *A*_*e*_, which should most robustly represent the behavior of pandemic strains that cause large numbers of cases in both old and young age groups. To obtain these values, we set *H*_*m*_ at 0.245 (the midpoint value from our main analyses; see Table 1), constrained *A*_*c*_ and *A*_*e*_ to be identical for H5N1 and H7N9, and estimated the parameter values by maximizing the constrained likelihood.

One hundred simulations were run for each pandemic scenario. To appropriately represent all sources of uncertainty, each simulation was run with independent values of *H*_*m*_, *A*_*c*_, and *A*_*e*_ drawn from the relevant sampling distributions.

Projections of *R*_*eff*_ (Fig. 3B, Fig. S8) were created for the UK, China, and Vietnam, assuming that a group 1 HA had constant seasonal dominance of 0%, 25%, 75%, or 100% from 2015 to 2060. For a given country, seasonal dominance scenario, and year, the fraction of each birth-year cohort with first exposure to each HA group was calculated as described in Methods (Reconstructing immune imprinting patterns) and used in conjunction with *H*_*m*_ to calculate the proportion of each birth-year cohort that would experience a ‘protected’ versus ‘unprotected’ course of infection with H5 or H7 IAV. The next generation matrix for a fully naïve population **M** (described in the previous section) was modified to separate ‘protected’ and ‘unprotected’ individuals, and account for the reduced infectiousness of ‘protected’ individuals (governed by *α*), and *R*_*eff*_ was calculated as the dominant eigenvalue of this matrix.

## Acknowledgements

We thank the Lloyd-Smith lab and the Worobey lab for helpful comments, C. Viboud for providing insight into historic influenza data, T. Mega and S. Wu for assistance compiling data, B. Cowling for sharing poultry exposure data, and P. Horby for sharing Vietnam contact data. K.G. is supported by the National Institute of General Medical Sciences of the National Institutes of Health (T32GM008185). M.A. is supported by the National Science Foundation Graduate Research Fellowship (DGE-1144087). M.W. is supported by the David and Lucile Packard Foundation. J.O.L-S. is supported by the National Science Foundation (EF-0928690), the Research and Policy for Infectious Disease Dynamics (RAPIDD) program of the Science and Technology Directorate, Department of Homeland Security, and Fogarty International Center, National Institutes of Health. The content is solely the responsibility of the authors and does not necessarily represent the official views of the National Institutes of Health. The authors declare no competing financial interests. Correspondence and requests for materials should be addressed to.

## Supplemental Information

### Supplementary Figures and Tables

**Fig. S1.**
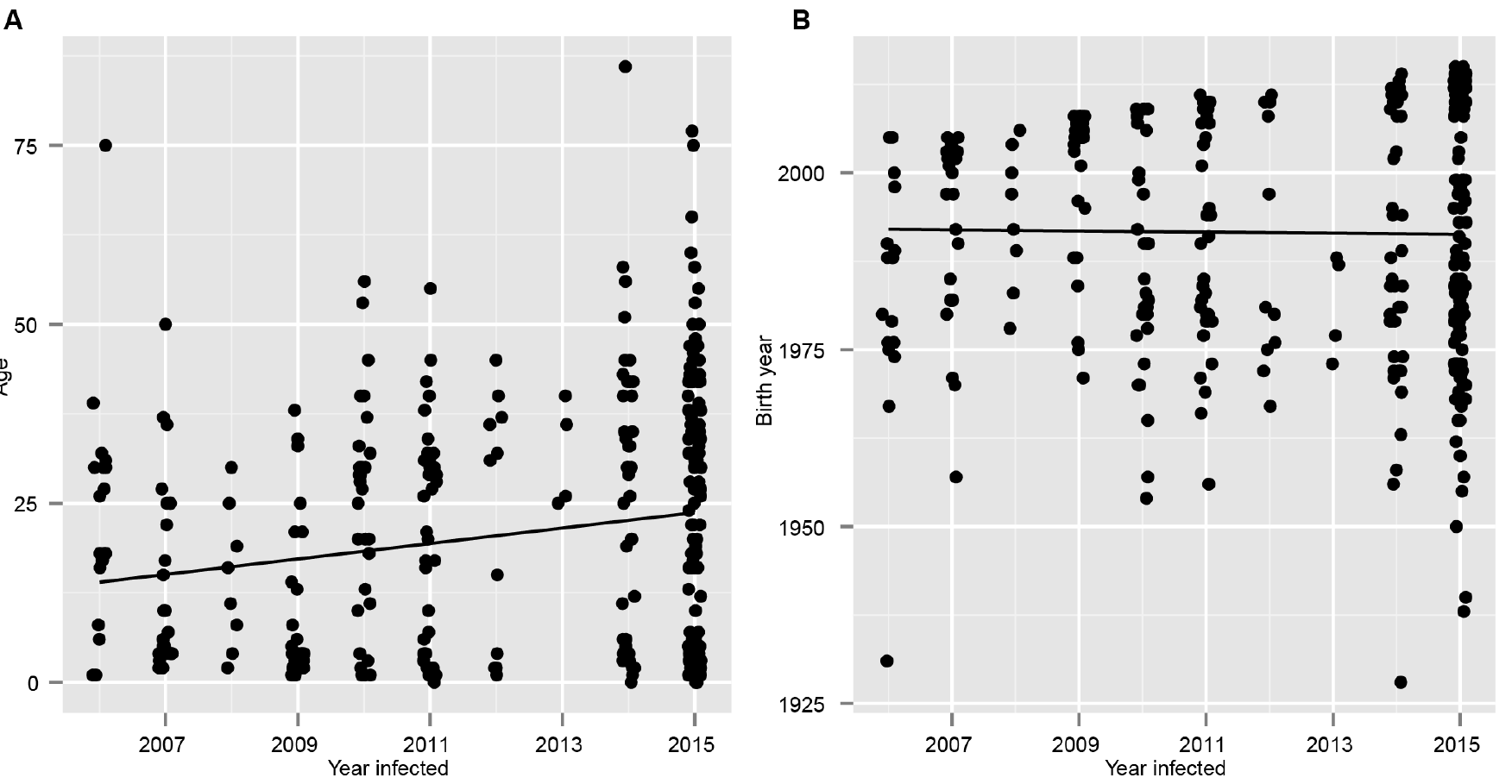
Trends in age and birth year of H5N1 cases over time. For 361 H5N1 cases observed in Egypt from 2006-2015, (**A**) Spearman’s rank correlation showed a significant positive association (p = 0.0003, one-sided test) between patient age and year of case observation. Points are jittered around the case observation year, and a least-squares trend line is shown. (**B**) Using the same test, no significant association was found for patient birth year. In support of the HA imprinting hypothesis, these results show that birth year is a more consistent predictor of severe infection risk than age-specific risk factors.

**Fig. S2.**
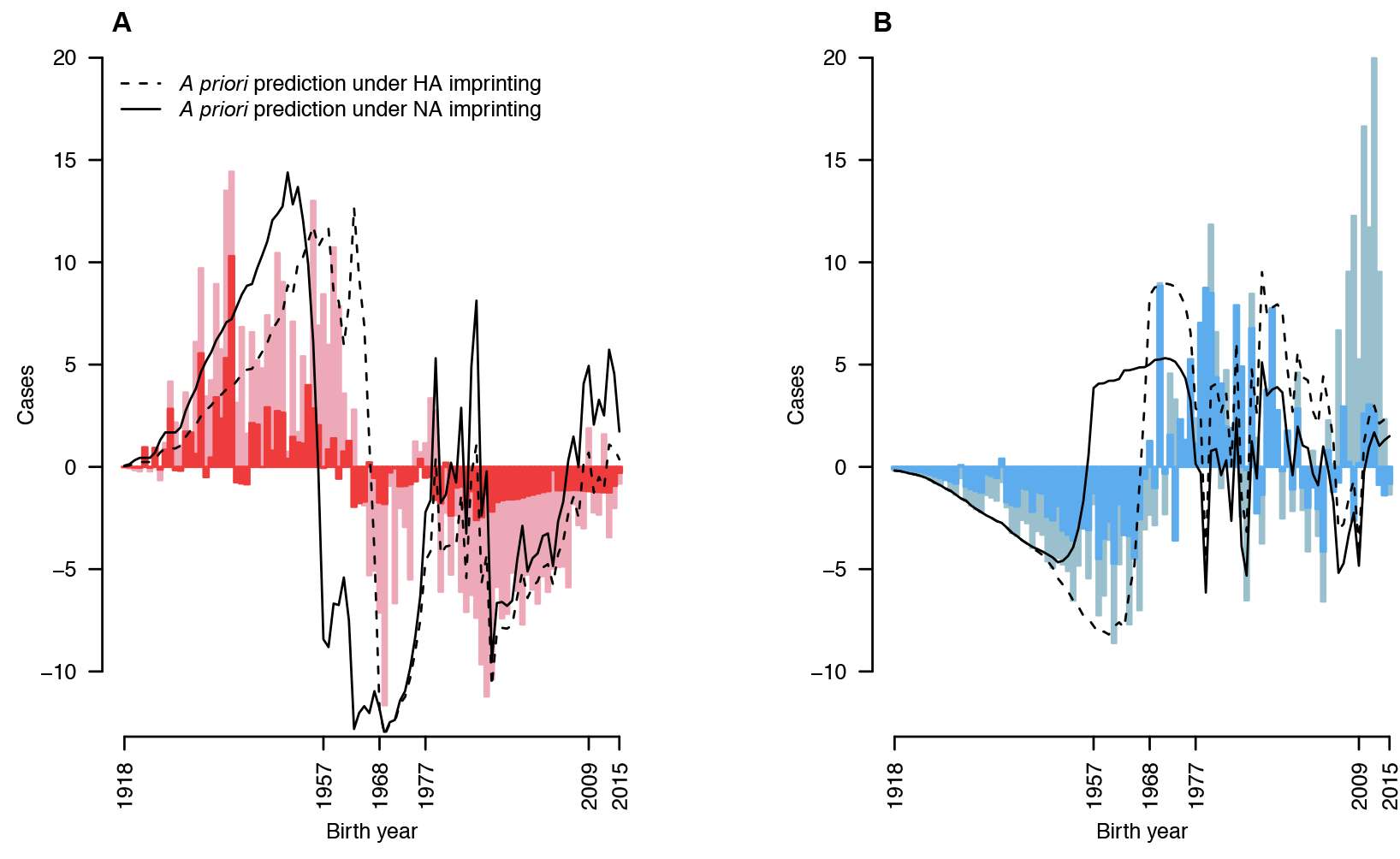
Comparison of NA and HA imprinting hypotheses to data. (**A**) H7N9. (**B**) H5N1. Bars show incidence (light colors) and mortality (dark colors) normalized to demography as in Figure 2C, Figure 2D. Overlaid lines show the *a priori* prediction based on HA imprinting (dashed line) or NA imprinting (solid line). During the period from 1957 to 1968 the NA imprinting prediction clearly fails to match observed incidence of excess severe infection or death from H7N9 or H5N1. Moreover, our modeling analysis indicates HA imprinting, not NA imprinting, is the dominant effect driving H5N1 and H7N9 severity patterns (see Table S2).

**Fig. S3.**
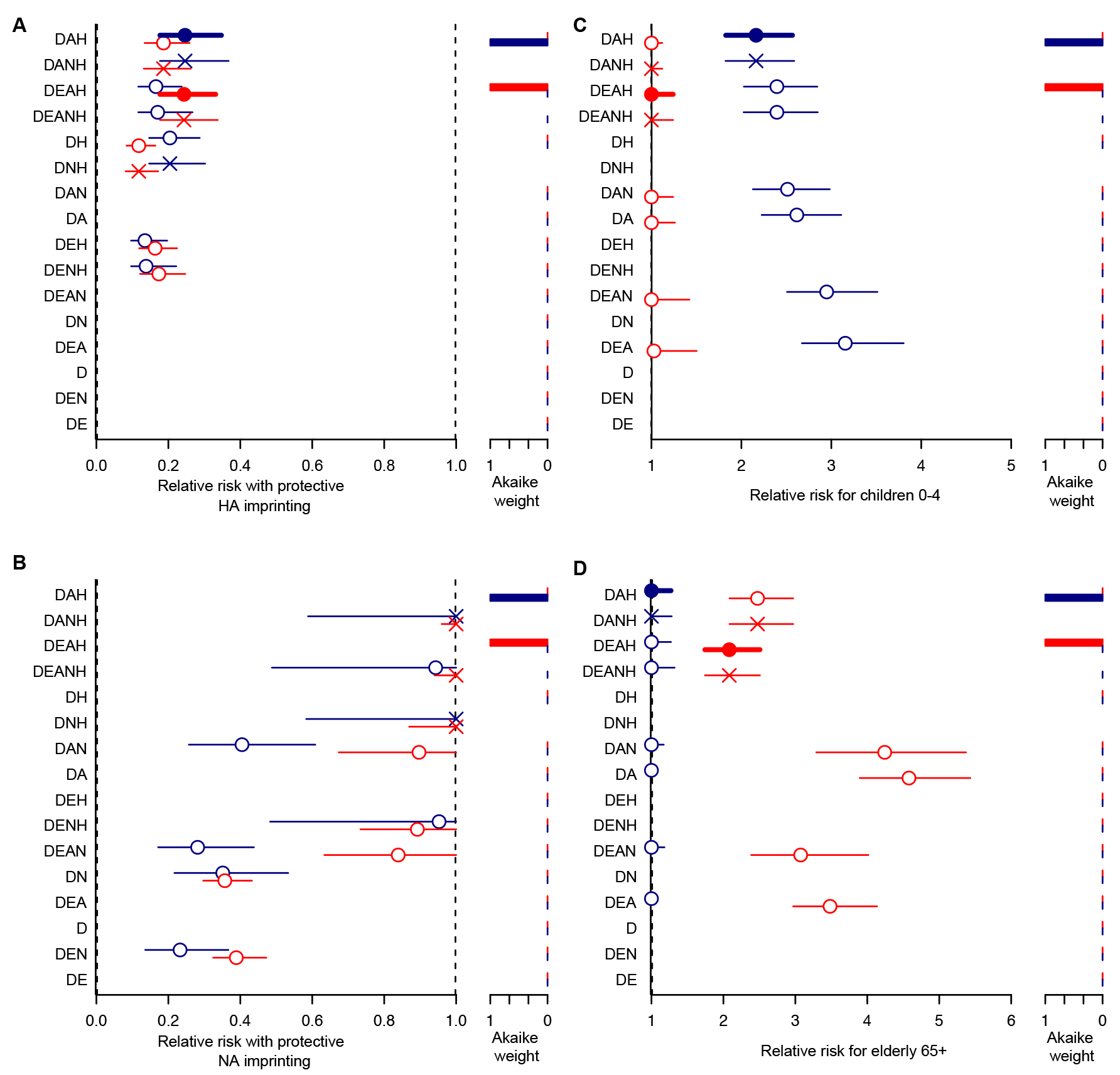
Parameter estimates for all models of H5N1 and H7N9 incidence. Maximum likelihood estimates and 95% likelihood profile CIs for parameters (**A**) *H*_*m*_ (**B**) *N*_*m*_ (**C**) *A*_*c*_ and (**D**) *A*_*e*_, as fit to H5N1 (blue) and H7N9 (red) incidence data. Parameters and model abbreviations are described in Table S1, and Methods. Demography, D, is included in all models. Additional tested factors include poultry exposure risk, E, age-based risk of severe morbidity in children and the elderly, A, NA imprinting, N, and HA imprinting, H. Dashed lines indicate boundary values imposed on parameters. Models are listed from top to bottom in order of decreasing model support, as fit to H5N1 data. Bar plots at right show Akaike weights, and the best model is shown with a filled symbol and bold line; all preferred models have definitive statistical support with Akaike weights > 0.99 (Table S2). In some cases, the MLE for parameter *N*_*m*_ was 1, indicating no NA imprinting effect, and the model with added factor N was identical to the model containing all the same factors except N. These degenerate models are represented with X’s, and were excluded from Akaike weight calculations.

**Fig. S4.**
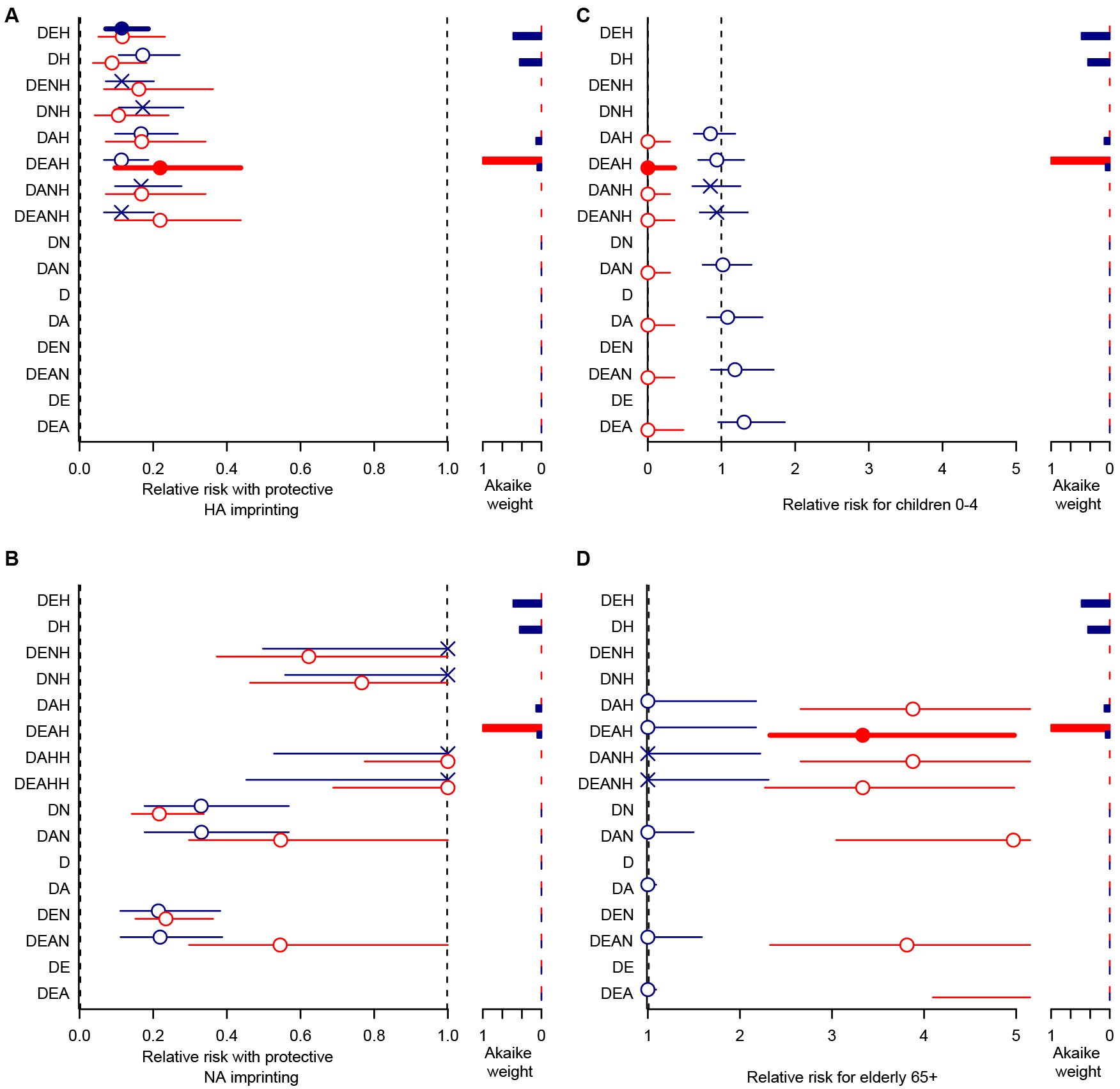
Parameter estimates for all models of H5N1 and H7N9 mortality. Maximum likelihood estimates and 95% likelihood profile CIs for parameters (**A**) *H*_*m*_ (**B**) *N*_*m*_ (**C**) *A*_*c*_ and (**D**) *A*_*e*_ as fit to H5N1 (blue) and H7N9 (red) mortality data. Parameters and model abbreviations are described in Table S1, and Methods. Models are listed from top to bottom in order of decreasing model support, as fit to H5N1 data. Dashed lines indicate boundary values imposed on parameters. Demography, D, is included in all models. Additional model factors include poultry exposure risk, E, age-based risk of severe morbidity in children and the elderly, A, NA imprinting, N, and HA imprinting, H. Bar plots at right show the Akaike weights for the models. The best model is shown with a filled symbol and bold line. X’s represent estimates from degenerate models, which were excluded from Akaike weight calculations. For H7N9, which is less pathogenic in birds and humans than H5N1, DEAH is the definitive preferred model for both infection and mortality, with Akaike weights > 0.99 in both cases. However, for H5N1, which is more pathogenic in birds and humans, no single mortality model is definitively preferred. Rather, we found some statistical support (Akaike weights > 0.05) for all mortality models that include HA imprinting (Table S2).

**Fig. S5.**
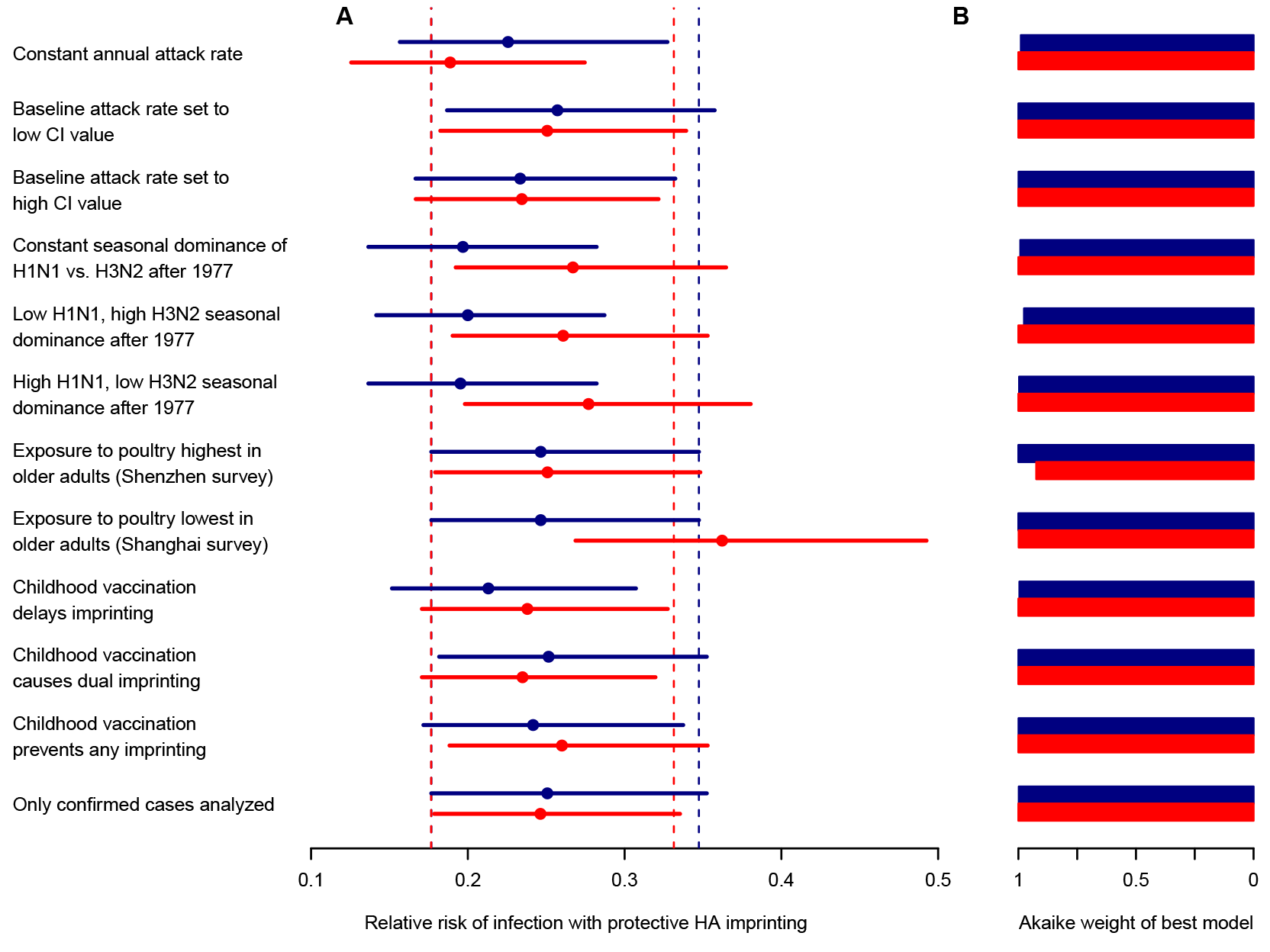
Sensitivity analyses. We tested the robustness of *H*_*m*_ parameter estimates (Table S1) and model selection results for nine variations on our standard model formulation (see Methods for details). (**A**) Maximum likelihood estimates and 95% likelihood profile CIs for parameter *H*_*m*_ as fit to incidence data using the best incidence model for each subtype (DAH for H5N1, in blue, and DEAH for H7N9, in red). Dashed lines show 95% CIs on *H*_*m*_ estimates from the preferred model, as presented in the main text analysis. (**B**) Akaike weights for models DAH (H5N1, blue) and DEAH (H7N9, red). These models remain strongly supported throughout all sensitivity analyses, with Akaike weights > 0.98 in all cases but one. The lowest Akaike weight was 0.92.

**Fig. S6.**
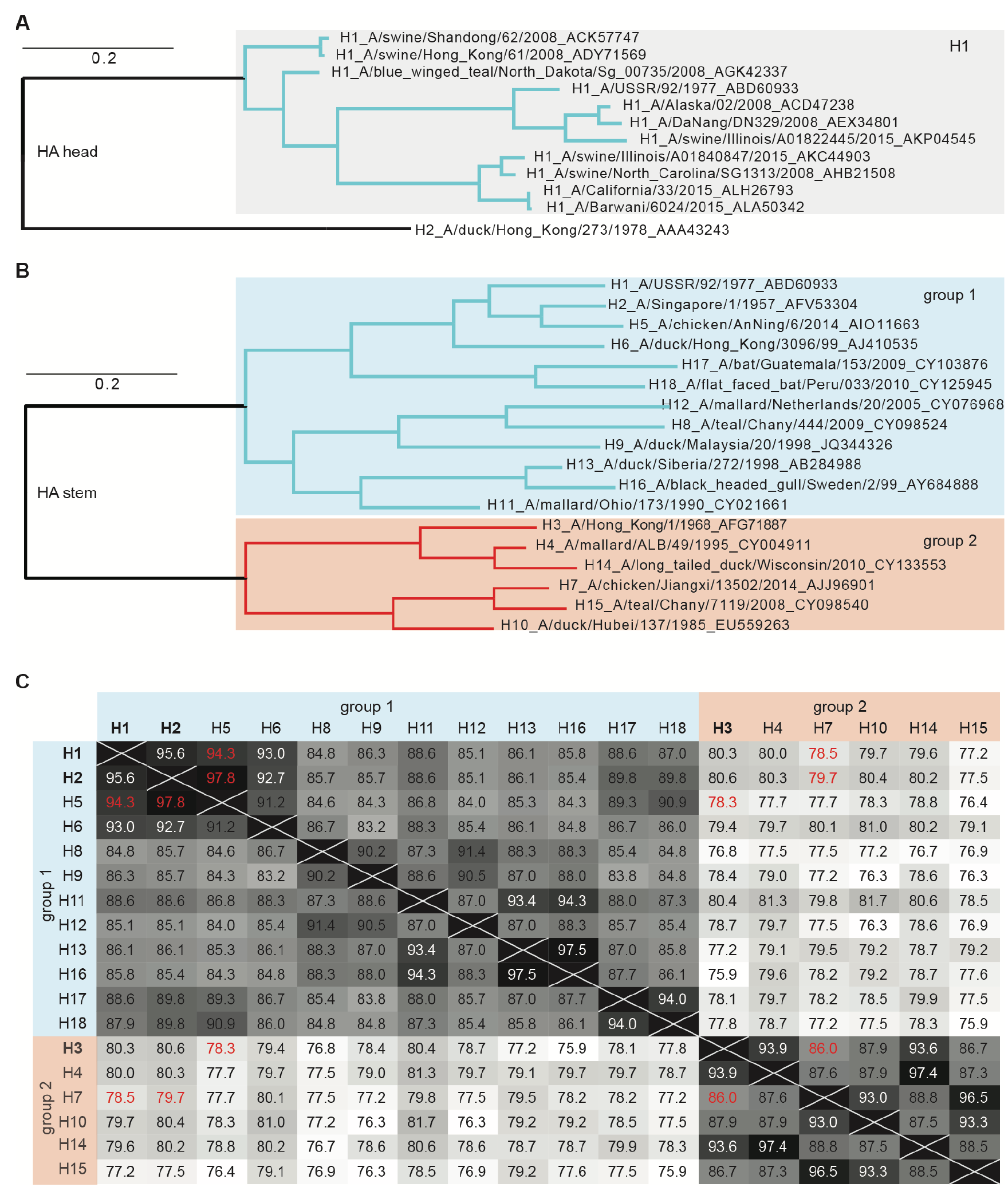
Phylogenetic and sequence analyses of HA amino acids. (**A**) Maximum likelihood phylogenetic tree of H1 HA globular head amino acid sequences. (**B**) HA stem domain amino acid tree. Both trees are drawn at the same scale. (**C**) Heat map showing pairwise similarities between HA stem domain amino acid sequences (same strains as in plot **B**). The number in each cell is the percent similarity (BLOSUM62 cost matrix) for the relevant pair of sequences, with darker colors indicating higher similarity. Pairwise comparisons for H5 and H7 versus H1, H2, and H3 (the variants that have circulated in human populations since 1918) are depicted in red. Each HA subtype’s stem domain is more similar to the other subtypes within its group than to any subtype in the opposing group, consistent with the observed imprinting pattern operating within, but not between, HA groups.

**Fig. S7.**
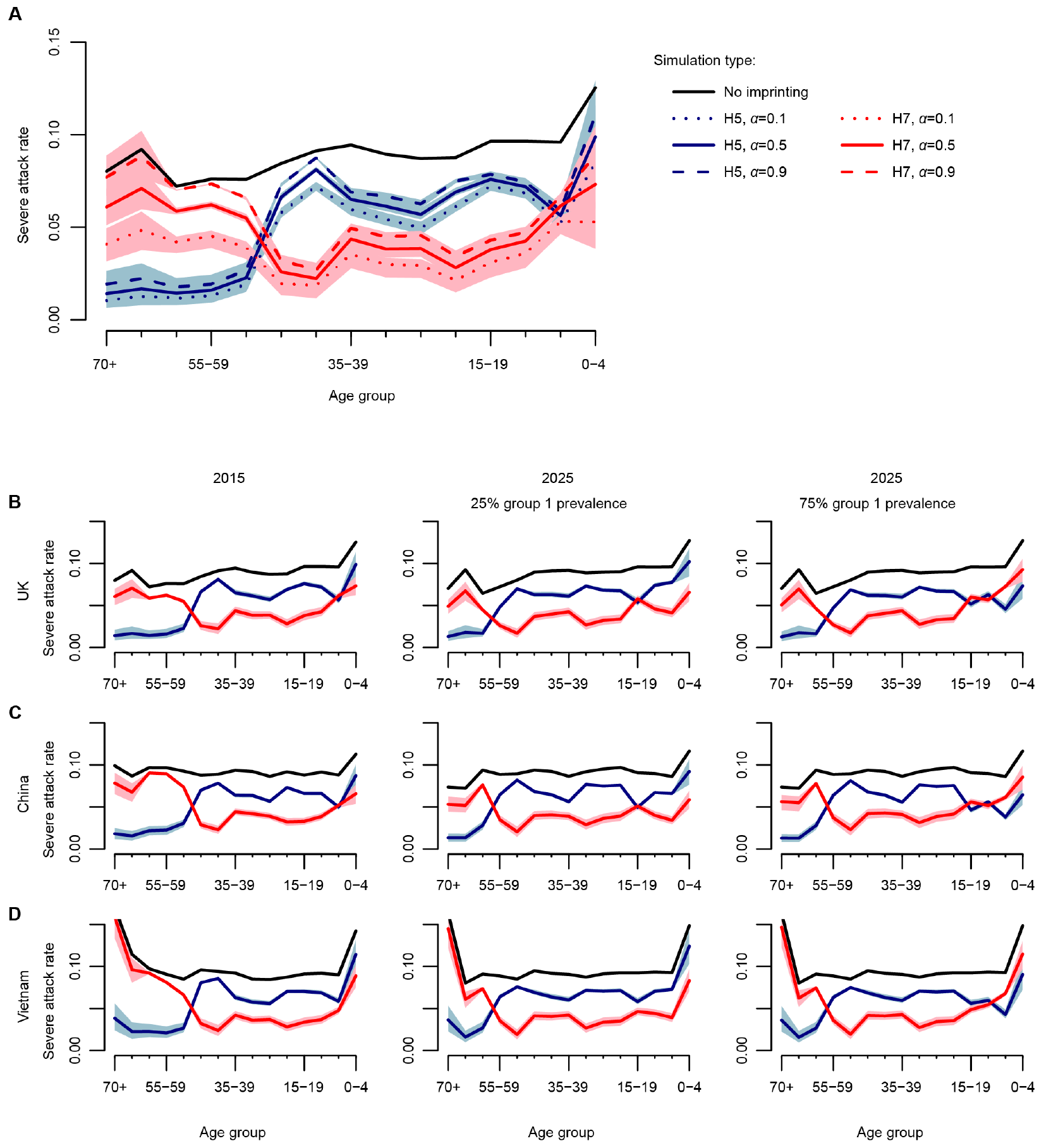
Projected age-structured severe attack rates during hypothetical future H5 (blue) and H7 (red) IAV pandemics. Simulations assume country-specific demography and age-structured mixing, HA imprinting, and age-based risk groups (Supplementary Text). Colored lines show the average, and shaded regions include 95% of 100 simulated outcomes. For comparison, the severe attack rate in an IAV pandemic with no HA imprinting is also shown (black). All simulations use *R*_0_ = 2.5. (**A**) Explores how changing the infectiousness of partially protected individuals (*α*), relative to unprotected individuals, affects the age-structured severe attack rate in a 2015 UK IAV pandemic. (**B-D**) Shows the results for three countries in 2015 and 2025 when *α* = 0.5. Two scenarios are considered for seasonal influenza circulation between 2015 and 2025: 25% group 1 AIV and 75% group 1 AIV.

**Fig. S8.**
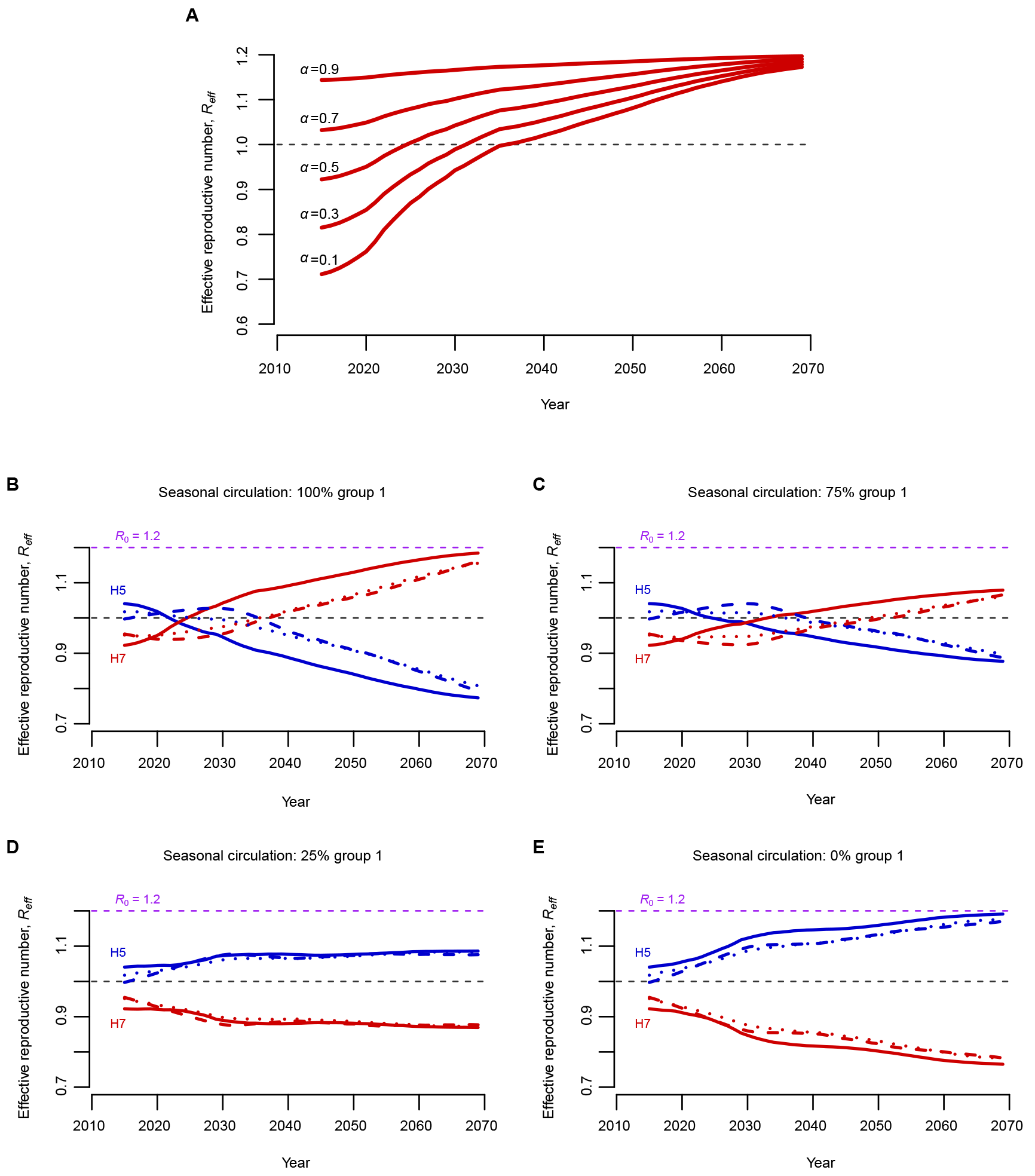
Projections of *R*_*eff*_ through time. (**A**) Projections of *R*_*eff*_ for an H7 IAV with *R*_0_ = 1.2 under different assumptions regarding the infectiousness of protected individuals (α). Projections are in the UK assuming 100% group 1 seasonal circulation after 2015. (**B-E**) Projections of how the *Rf* for a hypothetical H5 (blue) and H7 (red) IAV with *R*_0_ = 1.2 and *α* = 0.5 would change through time in the UK (solid line), China (dashed line), and Vietnam (dotted line) under four different post-2015 seasonal group 1 IAV circulation scenarios. Although general trends are consistent across countries, different demographic age structures and mixing patterns result in notable differences in the exact *R*_*eff*_ trajectory of each country.

**Table S1.**
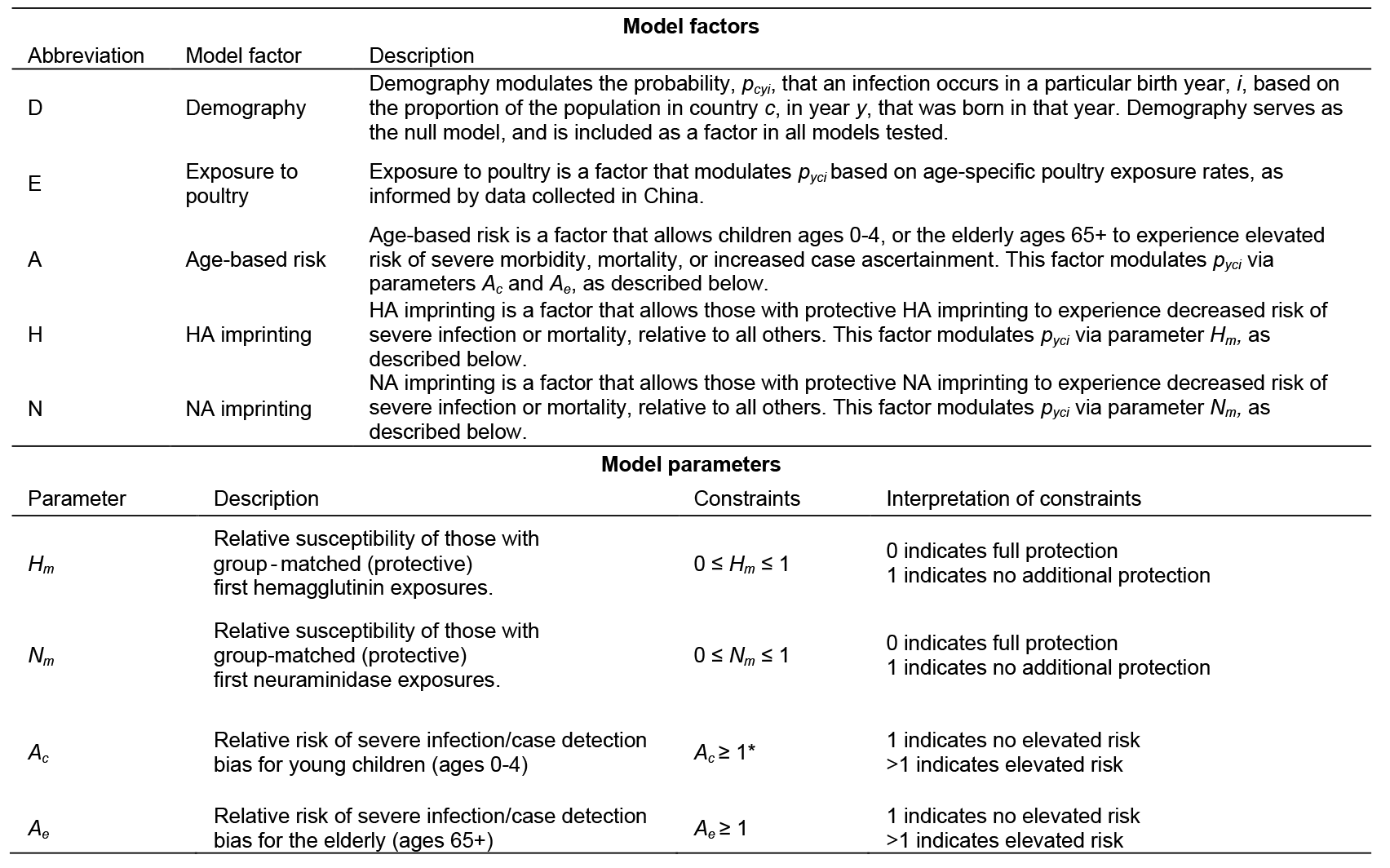
Summary of model factors and free parameters.

**Table S2.**
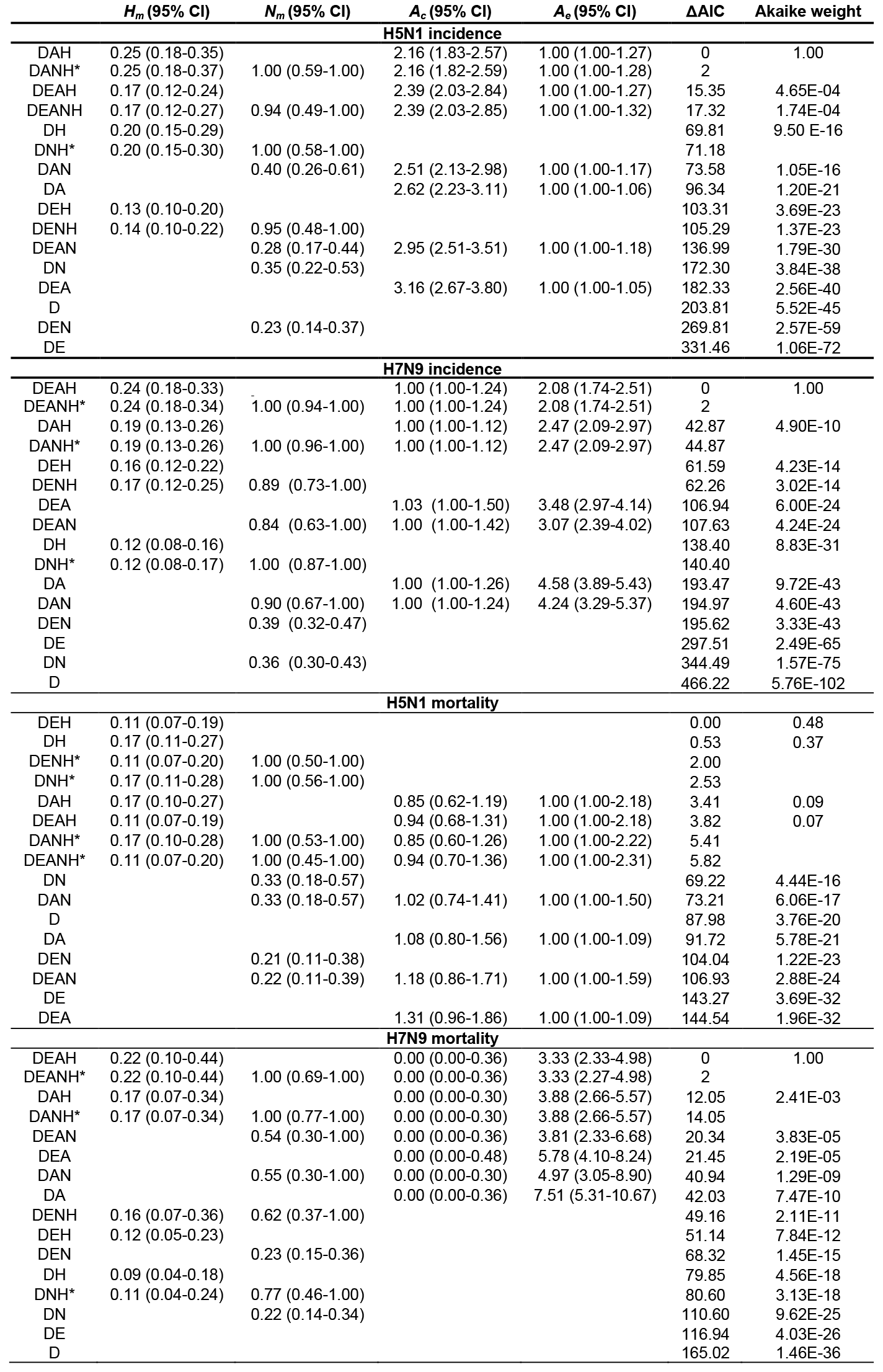
Incidence and mortality model results.

## Supplementary Text

### Contents

1. Likelihood functions
2. Model equations
3. Reconstructing immune imprinting patterns with vaccination
4. Annual intensity of influenza circulation from 1918-2015
5. Binomial exact test for mortality data
6. Analysis of novel subtypes other than H5N1 and H7N9
7. Parameter identifiability

### 1. <u>Likelihood functions</u>

The probability density function for the multinomial distribution is:

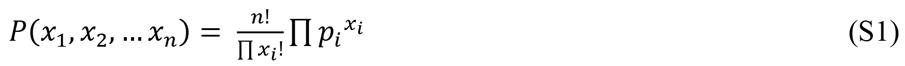

In our analysis, for a given country in a given year, we considered the multinomial probability of observing a certain number of cases or fatalities, *x*_*i*_, in each birth year, *i*. Each of our models assumes different factors may influence *p*_*i*_, the probability that any case has birth year *i*.

To find the likelihood of the full data, which comes from several countries in several case observation years, we multiply the multinomial probabilities from all relevant countries, *c*, and all relevant years of case observation, *y*. The full likelihood is:

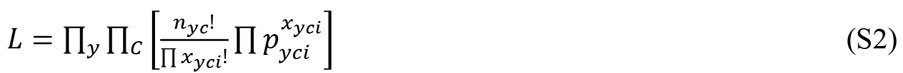

From equation S2, we obtain the full log likelihood:

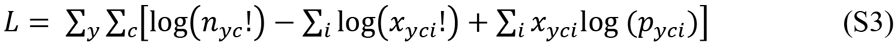

Maximum likelihood estimation was performed in R (version 3.2.0) and the optim() function was used to minimize the negative log likelihoods of candidate models. Code for model fitting is provided as a supplementary data file.

### 2. <u>Model equations</u>

Each model below assumes that a unique combination of five possible factors determine *p*_*yci*_ the probability that an infection or death observed in year*y*, country *c*, occurred in birth cohort *i* (Methods, Table S1). Demography (D) serves as the null hypothesis for the distribution of cases across birth years, and appears in every model. We assume all additional factors act independently. In all models, we normalize the probabilities to ensure 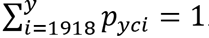.

#### 2.1 <u>Demography (D)</u>

This model assumes incidence in each birth year should be proportional to the fraction of the population of country *c*, year *y*, born in the year of interest.

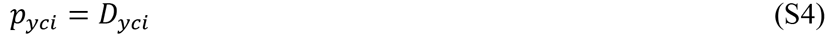

*D*_*yci*_ = Fraction born in year *i*, of the total population of country *c* born between 1918 and the case observation year (*y*).

#### 2.2 <u>Demography + exposure (DE)</u>

This model adds a factor describing poultry exposure risk across age groups.

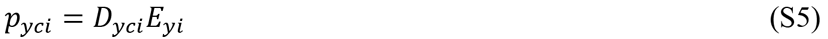

*E*_*yi*_ = Proportional risk in each birth year cohort, based on the frequency with which individuals of a given age (birth year *i*, in year *y*) contact poultry.

#### 2.3 <u>Demography + age-based risk (DA)</u>

This model introduces parameters that allow children ages 0-4 or the elderly ages 65+ to experience increased risk relative to other children and adults.

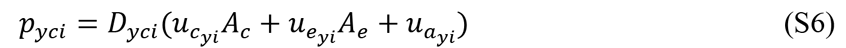

##### Group indicators

- *u*_*c*_*yi*__ = Indicator of membership in the age group representing high-risk children, taking value 1 for birth year cohorts of age 0-4 in year *y*, and 0 for all others.
- *u*_*a*_*yi*__ = Indicator of membership in the reference age group, taking value 1 for birth year cohorts of age 5-64 in year *y*, and 0 for all others.
- *u*_*e*_*yi*__ = Indicator of membership in the high-risk, elderly age group, taking value 1 for birth year cohorts of age 65+ in year *y*, and 0 for all others.

##### Parameters

- *A*_*c*_ = Proportional increase in risk for children ages 0-4, relative to the reference age group.
- *A*_*e*_ = Proportional increase in risk for the elderly, ages 65+, relative to the reference age group.

#### 2.4 <u>Demography + hemagglutinin imprinting (DH)</u>

This model introduces parameter *H*_*m*_, which allows individuals with group-matched first hemagglutinin exposures to experience decreased risk relative to all others.

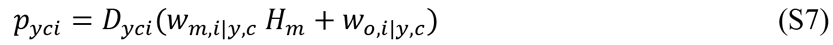

##### Group indicators

- *w*_*m,i|y,c*_ = Faction of each birth cohort with protective HA imprinting (group-matched to the challenge strain). This group should experience reduced risk under the HA imprinting hypothesis.
- *w*_*o,i|y,c*_ = Fraction of each birth cohort without protective HA imprinting. This group includes both IAV-nai’ve children and all individuals with HA-mismatched first exposures.

##### Parameters

- *H*_*m*_ = Proportional risk for individuals with group-matched first hemagglutinin exposures, relative to all others. *H*_*m*_ takes values in the range [0, 1], where 0 indicates full protection and 1 indicates no additional protection relative to the reference group.

#### 2.5 <u>Demography + neuraminidase imprinting (DN)</u>

This model introduces parameter *N*_*m*_, which, similarly to the DH model above, allows those with group-matched first neuraminidase exposures to experience decreased risk.

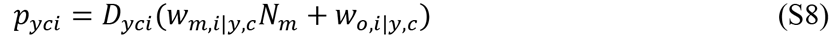

##### Group indicators

- *w*_*m,i|y,c*_ = Fraction of each birth cohort with NA imprinting group-matched to the challenge strain.
- *w*_*o,i|y,c*_ = Fraction of each birth cohort without protective NA imprinting.

##### Parameters

- *N*_*m*_ = proportional reduction in risk for individuals with group-matched first hemagglutinin exposures, relative to all others. *N*_*m*_ takes values in the range [0, 1], where 0 indicates full protection and 1 indicates no additional protection relative to the reference group.

#### 2.6 <u>Demography + neuraminidase imprinting + hemagglutinin imprinting (DNH)</u>

This model includes protection from first exposure to a group-matched HA (as in the DH model) as well as a group-matched NA (as in the DN model).

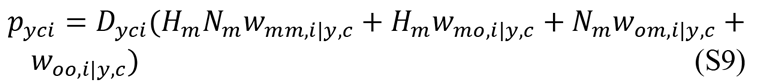

##### Group indicators

- *w*_*mm,i|y,c*_ = Fraction of each birth cohort with matched imprinting to HA and NA.
- *w*_*mo,i|y,c*_ = Fraction of each birth cohort with matched HA imprinting and mismatched NA imprinting.
- *w*_*om,i|y,c*_ = Fraction of each birth cohort with mismatched HA imprinting and matched NA imprinting.
- *w*_*oo,i|y,c*_ = Fraction of each birth cohort with mismatched imprinting to HA and NA, or that is naïve to IAV infection.

#### 2.7 <u>Demography + age-based risk + hemagglutinin imprinting (DAH)</u>

This model assumes that demography, age-based risk and HA imprinting determine overall risk.

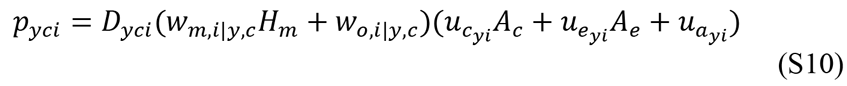

#### 2.8 <u>Demography + age-based risk + neuraminidase imprinting (DAN)</u>

This model assumes that demography, age-based risk and neuraminidase imprinting, determine overall risk.

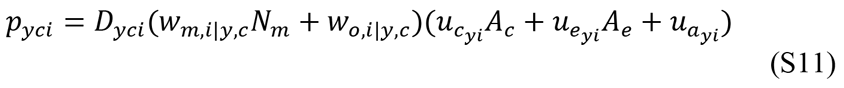

#### 2.9 <u>Demography + age-based risk + neuraminidase imprinting + hemagglutinin imprinting (DANH)</u>

This model assumes that demography, age-based risk, neuraminidase imprinting and hemagglutinin imprinting determine overall risk.

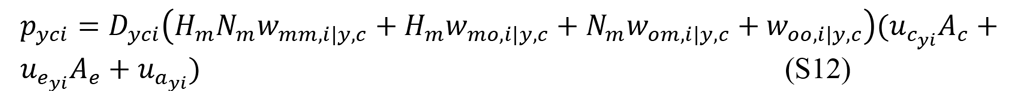

#### 2.10 <u>Demography + exposure + age-based risk (DEA)</u>

This model assumes that demography, exposure frequency and age-based risk determine overall risk.

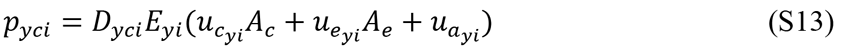

#### 2.11 <u>Demography + exposure + hemagglutinin imprinting (DEH)</u>

This model assumes that demography, exposure frequency and hemagglutinin history determine overall risk.

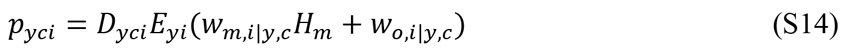

#### 2.12 <u>Demography + exposure + neuraminidase imprinting (DEN)</u>

This model assumes that demography, exposure frequency and neuraminidase history determine overall risk.

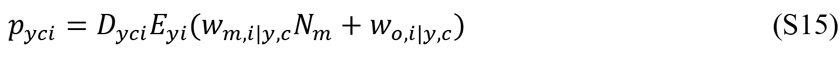

#### 2.13 <u>Demography + exposure + age-based risk + hemagglutinin imprinting (DEAH)</u>

This model assumes that demography, exposure frequency, age-based risk and hemagglutinin imprinting determine overall risk.

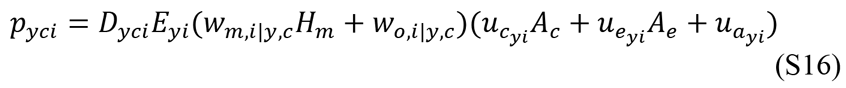

#### 2.14 <u>Demography + exposure + age-based risk + neuraminidase imprinting (DEAN)</u>

This model assumes that demography, exposure frequency, age-based risk and neuraminidase imprinting determine overall risk.

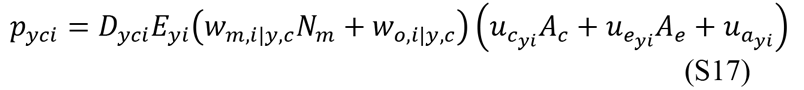

#### 2.15 <u>Demography + exposure + neuraminidase imprinting + hemagglutinin imprinting(DENH)</u>

This model assumes that demography, exposure frequency, hemagglutinin imprinting and neuraminidase imprinting determine overall risk.

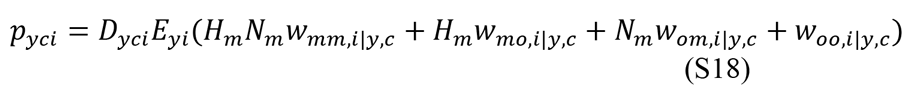

#### 2.16 <u>Demography + exposure + age-based risk + neuraminidase imprinting + hemagglutinin imprinting (DEANH)</u>

This model assumes that demography, exposure frequency, age-based risk, neuraminidase imprinting and hemagglutinin imprinting determine overall risk.

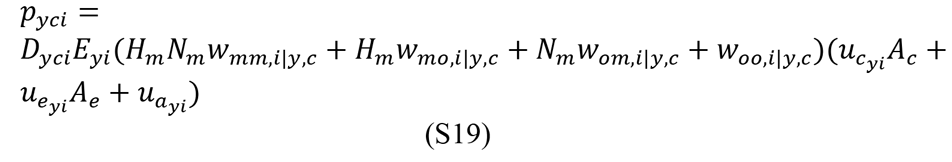

### 3. <u>Reconstructing immune imprinting patterns with vaccination</u>

When IAV-naïve children receive the seasonal influenza vaccine, the effect on HA imprinting in these children is unknown. We chose not to consider childhood vaccination in our main analysis, but instead to consider its possible effects via subsequent sensitivity analyses. Our rationale for omitting childhood vaccination in our main analysis was based on three main arguments.

First, childhood vaccination coverage is low in all H5N1-and H7N9-affected countries, so any effects of childhood vaccination on our results would be small. China and Thailand have recently launched public health initiatives to encourage early childhood influenza vaccination. However, even in these countries, recent estimates of childhood vaccine coverage are relatively low (and quite variable), ranging from 1% (*60*) to 30% (*61*) in Thailand, and from 26% (*62*) in China to <9% (*63*) in Hong Kong. For all other countries in our study, conservative estimates based on the number of vaccine doses purchased (*50*) show that even in the upper-bound limit that all vaccine doses were administered, and were distributed exclusively among children ages 0-9 years, childhood coverage would remain well below 5% in these countries (details below).

Second, early childhood influenza vaccination has only been widely recommended since the mid-late 2000’s (*61, 64*). Thus, the great majority of birth cohorts in our study would not have been affected at all. Since our study’s conclusions are driven most strongly by the dramatic change from group 1 to group 2 HA imprinting around the 1968 birth year, our conclusions are robust to variation in HA imprinting patterns in the very young birth cohorts that could have been affected by very recent, moderate increases in early childhood vaccination.

Third, for naïve children, single-dose influenza vaccine efficacy is exceptionally low, so the coverage estimates stated above strongly overestimate the effective vaccine coverage levels relevant to our analysis. IAV-naïve individuals require a series of two vaccine doses for effective protection (*64, 65*). However, in the United States, CDC data shows that only about 60% of children aged 6-24 months complete the two-dose course. We have not found equivalent data for H5N1-or H7N9-affected countries, but two-dose compliance is unlikely to be drastically better than in the US.

Given these arguments, we expected that childhood vaccination would have a minimal impact on our main findings, no matter how vaccination of IAV-naïve children affects HA imprinting at the individual scale. To test this expectation, we performed sensitivity analyses that considered three possible effects of imprinting: 1, vaccination of naïve children could prevent imprinting to either HA group, 2, vaccination of naïve children could cause dual imprinting to both HA groups or 3, vaccination of naïve children could delay the first natural infection and hence delay imprinting.

All of these sensitivity analyses required estimates of childhood vaccination coverage and efficacy for each country and year. Because reported estimates of childhood vaccine coverage vary considerably among studies, we estimated conservative, upper-bound limits on the true coverage to test the maximum effect childhood vaccination might have on our findings. We used data from Palache et al. (*50*) to determine the total number of vaccine doses distributed in each country of interest over time. For each year, we made the conservative assumption that all doses within the country were administered, and distributed uniformly among children ages 0-9. These estimates are conservative because in reality, not all doses are administered, and administered doses are actually distributed across all age groups, not exclusively in young children. Furthermore, while some naïve children would in reality receive two vaccine doses, we estimated upper-bound coverage levels as though all children required only one dose. We assumed vaccine efficacy was 60% in all non-pandemic years. This upper-bound efficacy estimate reflects that, as discussed above, at most 60% of naïve children complete the required two-dose course (*64*) This efficacy estimate is further conservative because even among children that receive two doses, the vaccine will not be perfectly protective due to antigenic drift.

In our first sensitivity analysis, we assumed that vaccination of IAV-naïve children delays the first natural infection, and hence delays imprinting, but does not affect imprinting otherwise. To estimate the fraction of each birth cohort with imprinting to particular seasonal subtypes, we incorporated the assumed vaccine-induced delay of imprinting by revising the definition of the annual attack rate on children, *a*_*j*_ (see equation 4 in Methods):

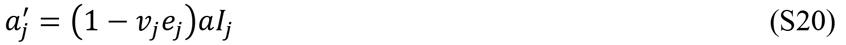

Here *V*_*j*_ describes childhood IAV coverage in year *j*, *e*_*j*_ describes IAV vaccine efficacy, *a* represents the baseline attack rate on children and *I*_*j*_ represents the year’s intensity score. With these modifications, we estimated vaccine-influenced imprinting patterns and performed model analyses as described in the Methods (Reconstructing immune imprinting patterns).

In our second and third sensitivity analyses, we assumed vaccination would prevent or replace imprinting in IAV-naïve children, so all children vaccinated before their first natural infection would either remain fully susceptible to both HA groups, or would simultaneously imprint to both groups. Here, it became necessary to keep track of the fraction first exposed via natural infection, the fraction first exposed via vaccination, and the fraction that remained naïve in the first 12 years after the birth year. We computed these using a recursive approach:

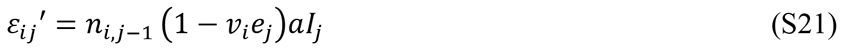

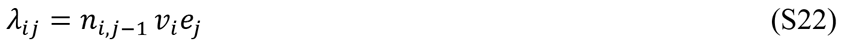

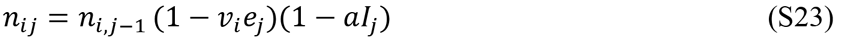

Here, as in equation 3 and equation 4, *ε*_*ij*_Ⅎ represents the fraction of birth cohort *i* that was first naturally infected in year *j* (prime indicates modification of the main text definition)*λ*_*ij*_. Misrepresents the fraction of birth cohort *i* that was naïve at the time of first vaccination in year *j. n_i_j* represents the fraction of cohort *i* that remained naïve at the beginning of year *j*, and was thus eligible for first infection or vaccination as year *j* progressed. In the first year of life (when *j* = i), *n*_*ii*_ is set to 1, indicating that all newborns are initially naïve. In all subsequent years, *j,* probabilities of first vaccination or natural infection were only applied to the naïve fraction of the cohort. Parameters *V*_*j*_, *e*_*j*_, *a* and *I*_*j*_ are as defined above.

After calculating relevant raw values of *ε*_*ij*_Ⅎ and *λ*_*ij*_, we applied a normalizing factor, *N*_*i|j*_Ⅎ to all *ε*_*ij*_Ⅎ and *λ*_*ij*_, As discussed in equation 4, the normalizing factor reflects the assumption that all individuals have their first natural infection or vaccination by age 12, and ensures that all relevant probabilities for a birth year sum to 1. It is given by:

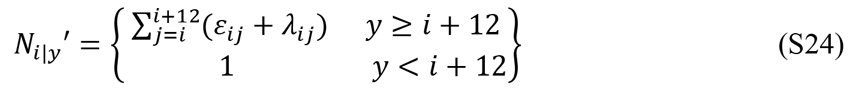

Finally, in the analysis assuming that vaccination of IAV-naïve children prevents imprinting, we always assigned the fraction of each cohort that was vaccinated before the first natural infection, **∑**_*i*_λ_*ij*_, to the mismatched-imprinting group, *w*_*o,i|y,c*_, as defined in equation 6 above. In the analysis assuming that vaccination of IAV-nai’ve children causes dual imprinting, we always assigned fraction **∑**_*i*_λ_*ij*_ to the matched-imprinting group, *w*_*m,i|y,c*_. Even at the upper limits of plausible vaccination coverage rates, our main results remained robust in all three sensitivity analyses (Fig. S5).

### 4. <u>Annual intensity of influenza circulation from 1918-2015</u>

Annual variation in IAV attack rate is most influential when estimating exposure weights for the post-1977 era of H3N2 and H1N1 co-circulation. During this time period, the dominant seasonal subtype can change annually, rather than on pandemic timescales. Conveniently, virological surveillance data from WHO collaborating laboratories are available from 1977-2015, providing a direct measure of annual IAV incidence. In years prior to 1977, the best estimates of the annual influenza burden come from pneumonia and influenza (P&I) excess mortality data (*66*). Thus, to estimate annual IAV intensity, we compiled estimates of incidence from virological surveillance data in years 1977-2015, and P&I excess mortality data in years 1918-1976.

Sensitivity analyses (Fig. S5) confirmed that our study’s results are quantitatively and qualitatively robust to uncertainty in the intensity index, and the results also remained robust when we replaced the intensity index with a constant value. Thus, we are confident that this index sufficiently captures broad patterns of historical IAV circulation intensity, and any assumptions do not unduly influence our study’s conclusions.

For 1997-2015 we obtained influenza surveillance data from our study’s six countries of interest. We defined raw annual incidence as: 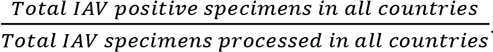. We specimens processed and the total positive specimens were reported. In years when the total number of reported specimens processed was less than 50 across all six countries of interest, (1997, 2000 and 2001), we substituted surveillance data from the United States. For 1977-1996 we were unable to find influenza surveillance data from countries affected by H5N1 and H7N9. Thus, we also substituted data from the United States for this time period (*49*).

After aggregating influenza surveillance data for all years from 1977-2015, we found that the observed annual incidence of influenza has increased predictably over time. A linear regression showed that the proportion of positive IAV specimens increased by about 0.0019/year (SE = 0.0006, p = 0.005). We assume this trend reflects a steady improvement in case detection efficiency, perhaps due to improved diagnostics or more targeted sampling, rather than a true increase in the annual influenza incidence proportion. Thus, we used this linear model to define yearly, expected incidence values for 1977-2015, and then defined annual circulation intensity as the ratio of observed to expected incidence.

For 1918-1976, we compiled published estimates of pneumonia and influenza (P&I) excess and expected mortality rates, per 100,000 population, for each year (*66–71*). Because northern hemisphere influenza seasons occur in the winter, estimates were not always reported on a calendar year basis. Instead, rates were often reported for a defined outbreak period, beginning and ending in specified months. In these cases, to adjust to the calendar year basis on which we defined birth cohorts, we first determined the fraction of outbreak months occurring within each calendar year, and then allocated the appropriate fraction of the season’s total excess mortality to either relevant year. Because we used our intensity time series to inform IAV imprinting patterns, we set excess mortality equal to zero in years dominated by type B influenza. In years when types A and B were co-dominant, we attributed half the total reported excess mortality to influenza A.

When verifying the comparability of excess mortality and incidence data, we found that the variance in post-1977 intensity estimates (informed by laboratory surveillance data) was greater than the variance in 1918-2015 estimates (informed by excess mortality data). To correct this discrepancy, we scaled the variance in the older data set to match the variance in more recent incidence estimates. We allowed a maximum intensity score of 2.5, to maintain a reasonable maximum annual attack rate of 0.75 or less. The maximum score applied only in years 1918, 1919, 1944 and 2009, all of which are recognized as years of intense influenza circulation.

### 5. <u>Binomial exact test for mortality</u>

One-sided binomial exact tests showed that census-excess H5N1 mortality was significantly less likely to occur in cohorts born before 1968 (estimated probability = 0.06, CI=0.00-0.19, p < 1e^−6^). For H7N9, census-excess mortality was significantly more probable in the same birth years (estimated probability = 0.94, CI=0.83-1.00, p < 1e^−7^). As introduced in the main text, the same test revealed similar patterns for the incidence of H5N1 (estimated probability = 0.00, CI=0.00-0.08, p < 1e^−10^) and H7N9 (estimated probability = 0.93, CI=0.84-1.00, p < 1e^−9^).

### 6. <u>Analysis of novel subtypes other than H5N1 and H7N9</u>

We searched the literature and avian influenza reports for clinically significant human cases of zoonotic IAV (other than H5N1 and H7N9) in which the year and country of the case was known and the age of the infected individual was reported (n=28). We excluded cases where the clinical manifestation was limited to conjunctivitis.

To examine whether the data were better explained by the Demography (D) or the Demography + Hemagglutinin imprinting (DH) model (described in sections 2.1 and Section 2.4 above), we performed a simple-versus-simple hypothesis test using the likelihood ratio as the test statistic. The null distribution of the test statistic (the distribution of likelihood ratio values generated under the assumption that the D model is true) was approximated using 250,000 simulated datasets. The quantile score of the true data’s likelihood ratio in this distribution was used to generate the p-value reported in the main text.

### 7. <u>Parameter identifiability</u>

A well-established pattern in influenza epidemiology is that young children and the elderly are at elevated risk for hospitalization and severe disease (*42–44*). Thus we designed our family of multinomial models to allow for elevated risk of severe disease in each of these age groups, via the parameters *A*_*c*_ and *A*_*e*_ respectively, to ensure that we accounted for all known age-related effects before testing our hypothesis of HA imprinting. However, a challenge arises in our study because there are very few H5N1 cases observed in the elderly, and few H7N9 cases observed in children (see Fig. 2A, Fig 2B). Thus, we have little statistical power to discern possible age-based risk effects in those groups. As a consequence, our results show elevated risk in age groups where we do see cases (i.e. in children for H5N1, and in the elderly for H7N9), but age-specific effects are not detected in age groups where cases do not frequently occur (Table S2). This leads to an effect where the age-based risk appears to act in concert with HA imprinting. It is thus prudent to consider the possibility that the age-based risk parameters (*A*_*c*_ and *A*_*e*_) and matched-imprinting protection parameter (*H*_*m*_) are not perfectly identifiable.

We performed three analyses to verify that our central findings are not meaningfully affected by this potential issue with parameter identifiability. First, we plotted twodimensional likelihood profiles for the parameter pairs of concern (figures not shown). These profiles show only weak positive correlations between the parameters *H*_*m*_ and *A*_*c*_ (for H5N1) and *H*_*m*_ and *A*_*e*_ (for H7N9). This indicates that identifiability is a minor concern. Second, from these profiles, we computed marginal 95% confidence intervals, and found values only slightly different from the univariate confidence intervals reported in Table 1 in the main text. For H5N1, the univariate CI on *A*_*c*_ was 1.83-2.57, and the marginal CI from the bivariate analysis was 1.75-2.65. For H7N9, the univariate CI on *A*_*e*_ was 1.74-2.51, and the marginal CI from the bivariate analysis was 1.68-2.58. Again, the small differences between univariate confidence intervals and marginal, bivariate confidence intervals indicate that identifiability is not a major issue for these parameters.

Third, as a final check, we combined H5N1 and H7N9 data into a single model, and constrained parameters *H*_*m*_, *A*_*c*_ and *A*_*e*_ to take the same values for both viruses (see details above in section 7). We then computed the maximum likelihood estimates for all three parameters, using the combined data set. The combined data set contained large numbers of cases in both old and young age groups, and gave us more power to estimate age-specific risk effects. The consensus estimates for *A*_*c*_ (1.78, 95% CI 1.53-2.07) and *A_e_* (1.91, 95% CI 1.64-2.23) were statistically indistinguishable from the estimates of *A*_*c*_ (using H5N1 data) and *A*_*e*_ (using H7N9 data) presented in the main text analysis.

